# Phosphorylation Determines Whether Neuroligin-3 is at Excitatory or Inhibitory Synapses in Different Regions of the Brain

**DOI:** 10.1101/2022.07.23.501257

**Authors:** Bekir Altas, Liam P. Tuffy, Annarita Patrizi, Kalina Dimova, Tolga Soykan, Andrea J. Romanowski, Colin D. Robertson, Saovleak N. Khim, Garrett W. Bunce, Mateusz C. Ambrozkiewicz, Oleksandr Yagensky, Dilja Krueger-Burg, Matthieu Hammer, He-Hsuan Hsiao, Pawel R. Laskowski, Lydia Dyck, Marco Sassoè-Pognetto, John J.E. Chua, Henning Urlaub, Olaf Jahn, Nils Brose, Alexandros Poulopoulos

## Abstract

Neuroligin-3 is a postsynaptic adhesion molecule involved in development, function, and pathologies of synapses in the brain. It is a genetic cause of autism and a potent component of the tumor microenvironment in gliomas. There are four Neuroligins that operate at distinct synapse types, selectively interacting with presynaptic adhesion and postsynaptic scaffold proteins. We investigated the subcellular localization and scaffold specificities of synaptic Neuroligin-3 and demonstrate an unexpected pattern of localization to excitatory synapses in cortical areas, and inhibitory synapses in subcortical areas. Using phosphoproteomics, we identify Neuroligin-3-specific serine phosphorylation in cortex and hippocampus that obstructs a key binding site for inhibitory synapse scaffolds. Using *in utero* CRISPR/Cas9 knockout and replacement with phosphomimetic mutants, we demonstrate that phosphorylation at this site determines excitatory versus inhibitory synapse localization of Neuroligin-3 *in vivo*. Our data reveal a mechanism that differentially regulates the balance of Neuroligin-3 between excitatory and inhibitory synapses, adding to our emerging understanding of their role in the development of brain connectivity and associated pathologies.

## INTRODUCTION

Neuroligins (NLs) are important cell adhesion molecules in the development of neural connectivity. Mutations in NL3 were among the first known monogenetic causes of non-syndromic autism in humans^1, 2^ and genetic models of autism in mice ^3–6^. NL3 also emerged as a critical component of the tumor microenvironment in the brain. NL3 shed extracellularly by synaptic activity promotes glioma spread^7^, while gliomas fail to grow in NL3 knockout mice^8^.

Among the four NL genes, mutations in NL3 cause the widest range of phenotypes at distinct synapse types depending on brain region^6, 9–13^. In contrast, other NLs are more consistently synapse-type selective. NL1 localizes and functions at excitatory synapses^14–16^, while NL2 localizes and functions at inhibitory synapses and subtypes of cholinergic and dopaminergic synapses^16–21^. NL4 in mice is restricted to subsets of inhibitory synapses, predominantly glycinergic^22–24^. The specificity of NL3 is reported mixed at excitatory and inhibitory synapses in culture^25^ and cerebellum^9^, and inhibitory in striatum ^21^ and retina^26^.

The absence of documented synapse-type specificity for NL3, as well as the range of synaptic defects reported in *Nlgn3* mutants, prompted us to systematically examine NL3 localization across brain regions. Using knockout-validated antibodies, we show that NL3 selectively associates with either excitatory or inhibitory synapses in cortical or subcortical brain areas, respectively. We went on to search for mechanisms that could regulate the differential localization of NL3 in different parts of the brain and identified that region-specific phosphorylation controls binding specificities for inhibitory versus excitatory postsynaptic scaffolds. Using an in utero CRISPR replacement strategy, we demonstrate that this phosphorylation-dependent binding mechanism determines the subcellular localization of NL3 *in vivo*.

## RESULTS

### Non-Specific dendritogenic and synaptogenic effects of NL3 overexpression in cultured neurons

We found that overexpression of HA-NL3 in cultured cortical neurons causes notable changes in neuron morphology, including increases in dendritic arborization and in the number of dendritic filopodia, as seen in previous studies involving exogenous expression of NLs^19, 27, 28^. These effects were elicited via the NL3 cytoplasmic domain, as overexpression of a truncated NL3 variant lacking the intracellular domain (HA-NL3^ΔICD^) did not produce these effects (Extended Data Fig. 1).

In addition, NL3 overexpression in cultured neurons caused pronounced effects on the density of presynaptic contacts (Extended Data Fig. 1A-1C), as previously demonstrated^15, 27, 29–31^. Overexpression of HA-NL3 increased the density of both excitatory and inhibitory presynaptic contacts as seen by immunolabeling for the corresponding presynaptic marker proteins VGluT1 and VIAAT (Extended Data Fig. 1B and 1C). The effects of NL3 overexpression on the recruitment of presynaptic terminals were sufficiently mediated by the NL3 extracellular domain alone, since ΔICD variants of NL3 were equally effective, similar to ΔICD effects seen in previous studies^32, 33^.

Overall, these data show that NL3 overexpression in cultured neurons elicits the same potent synaptogenic and dendritogenic effects that are commonly caused by overexpression of NL family members^15, 27, 29^, without specificity for excitatory or inhibitory synapses.

### Differential localization of NL3 to subsets of excitatory vs. inhibitory synapses in cortical vs. subcortical areas

We systematically examined the differential localization of endogenous NL3 to excitatory vs. inhibitory synapse types across the rodent brain with an antibody we previously generated^34^. To reliably determine NL3 localization, we first assessed antibody specificity for NL3 by comparing immunoreactivity in brain sections from wild type (WT) vs. *Nlgn3* knockout (NL3 KO) mice (Fig. 2A). We determined that our anti-NL3 antibody displayed minimal labeling on brains lacking NL3, confirming that the punctate immunolabeled signal in WT brains corresponds to endogenous NL3 clusters.

With this validated antibody, we examined the distribution of NL3 puncta across the adult mouse brain. Interestingly, NL3 puncta appeared in two distinct sizes depending on the brain region examined. Small NL3 puncta appear in cerebral cortex, hippocampus, and the molecular layer and deep nuclei of cerebellum. Large NL3 puncta were prevalent in olfactory bulb, basal ganglia, thalamus, brain stem, and the granule-cell layer of cerebellum (Table 1).

**Table 1.** Abundance, size, and synapse-type localization of NL3 puncta across brain regions. The left column lists brain regions (bold) and subregions included in the analysis. Column two tabulates the relative abundance and size of NL3 puncta. Abundance is indicated by 1 (low) to 3 (high) marks, * for small puncta and ♢ for large puncta. Lack of punctate immunoreactivity in a region is indicated by /. The third and fourth columns note the presence (+) or absence (-) of NL3 puncta colocalizing with Gephyrin or PSD95, respectively. Areas not examined are marked as n.a.

|  | NL3 | NL3/Geph | NL3/PSD95 |
| --- | --- | --- | --- |
| **Olfactory Bulb** |  |  |  |
| Glomerular layer | ◇ | + | - |
| External plexiform layer | ◇ | + | - |
| Mitral cell layer | / | - | - |
| Internal plexiform layer | ◇◇ | + | - |
| **Cerebral Cortex** | ** | - | + |
| **Hippocampus** |  |  |  |
| CA1, stratum pyramidale | ** | - | + |
| CA1, stratum oriens and radiatum | ** | - | + |
| CA1, stratum lacunosum-moleculare | ** | - | + |
| CA3, stratum pyramidale | ** | - | + |
| CA3, stratum oriens and radiatum | ** | - | + |
| **Dentate gyrus** |  |  |  |
| Granule cell layer | * | - | + |
| Molecular layer | *** | - | + |
| Hilus | ** | - | + |
| **Basal ganglia** |  |  |  |
| Caudate Putamen | ◇◇ | + | - |
| Globus pallidus | ◇◇◇ | + | - |
| **Thalamus** |  |  |  |
| Mediodorsal nucleus | ◇◇◇ | + | - |
| Centrolateral nucleus | ◇◇◇ | + | - |
| Paraventricular thalamic nucleus | ◇◇◇ | + | - |
| Reticular nucleus | ◇ | - | n.a. |
| Ventral lateral nucleus | ◇ | + | - |
| Rhomboid nucleus | ◇ | + | - |
| Reuniens nucleus | ◇ | + | - |
| Latero dorsal nucleus | ◇ | - | - |
| **Brainstem** | ◇◇◇ | + | - |
| **Cerebellum** |  |  |  |
| Molecular layer | ** | - | + |
| Purkinje cell layer | / | - | - |
| Granule cell layer | ◇◇◇ | + | + |
| Cerebellar nuclei | ** | - | n.a. |

We next investigated whether these puncta correspond to synaptic clusters of NL3 with excitatory or inhibitory synapse specificity. We co-labeled endogenous NL3 with excitatory and inhibitory postsynaptic markers PSD95 and Gephyrin, respectively. We found strong association of NL3 puncta with synaptic markers, indicating predominantly synaptic localization of NL3 clusters. Further, NL3 showed strong preference for either excitatory or inhibitory synapses that correlated with puncta size and varied between regions. In areas with small puncta, NL3 colocalized with PSD95 at excitatory postsynapses (Fig. 1B) with no appreciable overlap with Gephyrin at inhibitory postsynapses (e.g. cortex in Fig. 1D). In areas with large puncta, NL3 colocalized with Gephyrin (Fig. 1D) with no appreciable overlap with PSD95 (e.g. brainstem in Fig. 1B). In cerebellum, NL3 displayed layer-specific selectivity localizing to excitatory synapses in the molecular layer and deep nuclei, and to subsets of both excitatory and inhibitory synapses of cerebellar glomeruli in the granule cell layer (Table 1, Fig. 1, Extended Data Fig. 2), consistent with observations previously made in cerebellum using a distinct anti-NL3 antibody^9^. Together, our data show that endogenous NL3 displays strong synapse-type selectivity that is distinct across brain areas.

**Figure 1.**
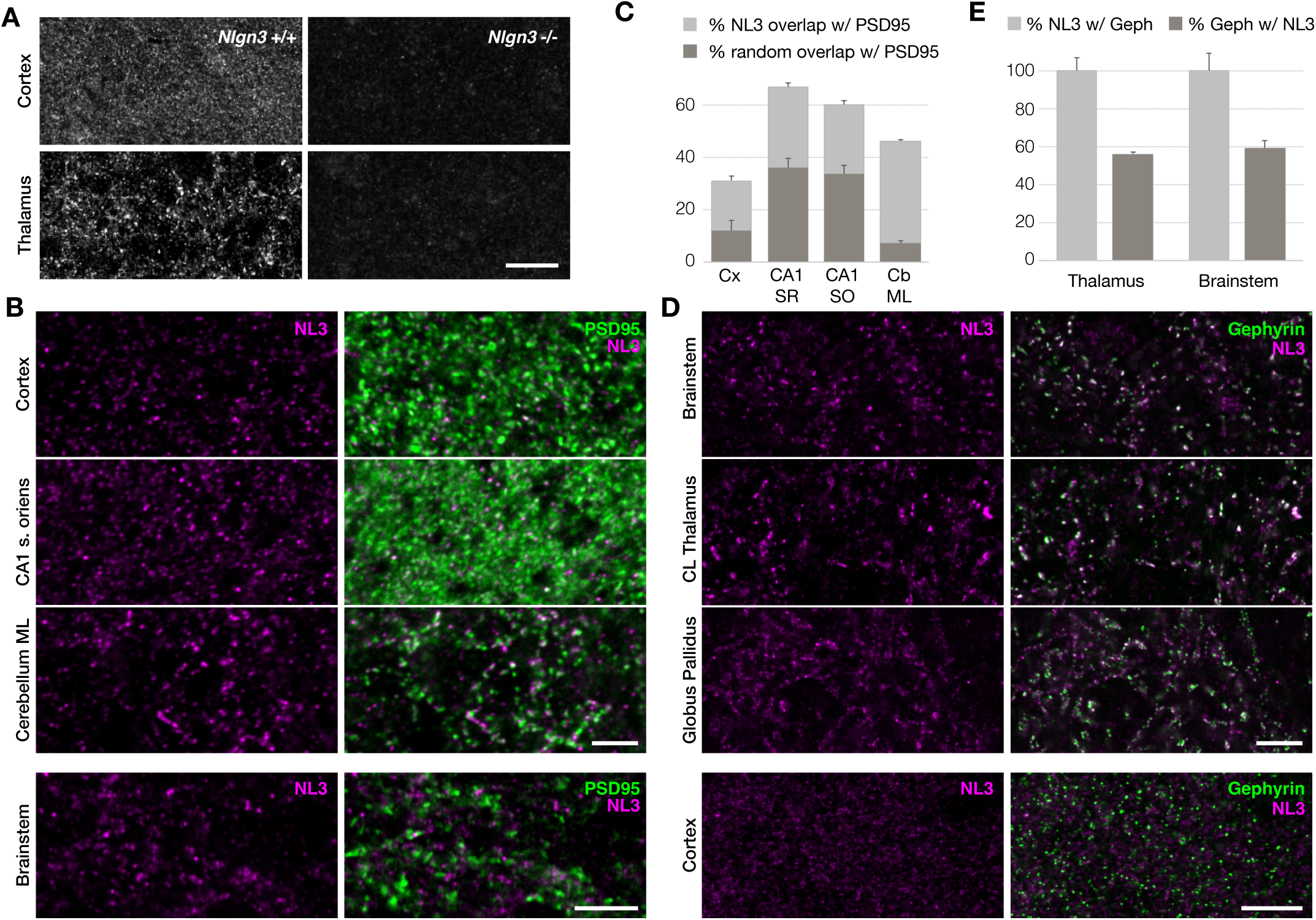
Endogenous NL3 localizes to excitatory synapses in cortical, and to inhibitory synapses in subcortical regions. (A) Validation of NL3 antibody immunolabeling in WT versus *Nlgn3* KO tissue shows NL3-specific punctate signal. Cortex displays small puncta and thalamus displays large puncta (scale bar, 45 μm). (B) NL3 colocalizes with subsets of PSD95 puncta in cerebral cortex, hippocampus CA1 stratum oriens, and cerebellum molecular layer (scale bar, 3 μm) but not in brainstem (scale bar, 14 μm). (C) Quantification of NL3 percentage overlapping with PSD95 puncta versus randomized overlap in cerebral cortex (Cx), hippocampus CA1 stratum radiatum (CA1 SR), stratum oriens (CA1 SO), and cerebellum molecular layer (CB ML). (D) NL3 significantly colocalizes with Gephyrin in brainstem, thalamus centrolateral nucleus, and globus pallidus, but not in cerebral cortex (scale bar, 7 μm). (E) Quantification of the percentage of NL3 that overlaps with Gephyrin, and the percentage of Gephyrin that overlaps with NL3 in thalamus and brainstem. NL3 is fully localized to a subset of inhibitory postsynapses in thalamus and brainstem.

Quantifications show that only a proportion of excitatory and inhibitory synapses contain NL3, indicating further synapse subtype specificities in all areas examined. Of total PSD95 puncta, overlap with NL3 reached 30% in cortex, 60-70% in hippocampus, and 45% in the molecular layer of the cerebellum (Fig. 1C). Correspondingly in olfactory bulb, striatum, thalamus, and brainstem, all NL3 puncta localize with Gephyrin, though only about half of total Gephyrin puncta contain NL3 (Fig. 1E). We examined whether NL3-containing inhibitory synapses segregated with either glycinergic or GABAergic subtypes, but found no such preference. Rather, NL3 was detected at GABA_A_R-positive, GlyR-positive, and mixed GABAergic-glycinergic postsynapses in brainstem (Extended Data Fig. 2).

These data reveal a general pattern of binary and mutually exclusive synapse-type specificity for NL3, and further selectivities –yet to be determined – for subsets of excitatory synapses in cortical areas and subsets of inhibitory synapses in subcortical areas. This robust and regulated synapse-type selectivity is peculiar to NL3 since other NLs show more consistent preferences for synapse types throughout the brain.

### Serine phosphorylation of the NL3 Gephyrin-binding site in cortical areas

The region-specific synapse-type selectivity of NL3 indicates the existence of a regulatory mechanism that determines NL3 association with excitatory or inhibitory postsynapses. The cytoplasmic domain of NL3 contains binding sites for both PSD95, the key scaffold protein of excitatory synapses, and for Gephyrin, the key scaffold protein of inhibitory synapses^14, 19, 35^ (Fig. 2A and 4). We hypothesized that these binding sites may be regulated by phosphorylation in a region-specific manner. To test this, we applied a targeted phospho-proteomic approach to identify potential phosphorylation of endogenous NL3 immunopurified from rat brain (Fig. 2B). Analysis by mass spectrometry (MS) detected a phosphopeptide that unequivocally represented a new NL3 phosphorylation site at serine 799 (S799), which is part of a conserved Gephyrin-binding consensus motif in the NL3 cytosolic domain (Fig. 2C). MS/MS sequencing of this monophosphorylated peptide yielded no evidence of phosphorylation on proximal residues, including tyrosine 792 (Y792), which corresponds to a known phosphorylation site in NL1^36^.

**Figure 2.**
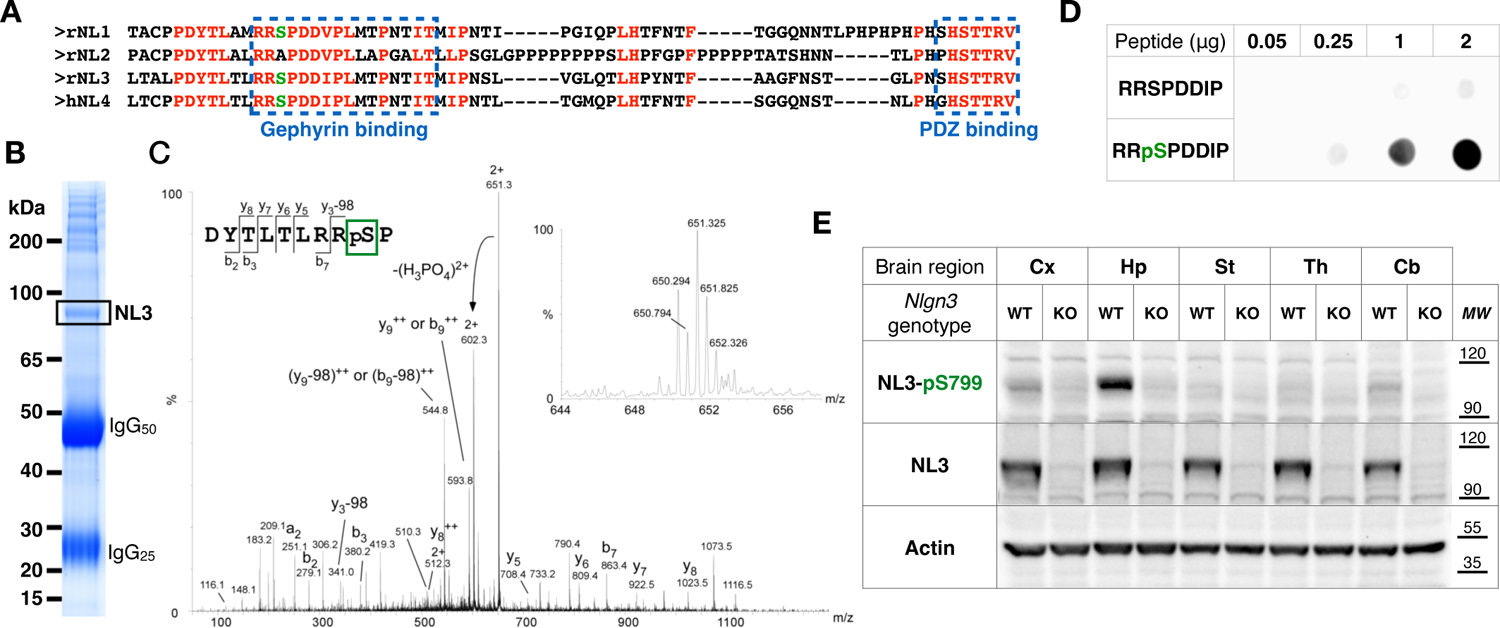
Identification of endogenous serine phosphorylation within the Gephyrin-binding site of NL3. (A) Alignment of the intracellular domains of rat NL1-NL3 and human NL4. Conserved and conservative residues are in red. PDZ- and Gephyrin-binding sites are indicated in blue. The serine within the Gephyrin-binding site corresponding to NL3-pS799 is highlighted in green. (B) Coomassie stained SDS-PAGE of anti-NL3 immunopurified eluate from rat brain extract. The extracted NL3 band as identified by mass spectrometry is boxed. Light (IgG_25_) and heavy (IgG_50_) immunoglobulin chains from the antibody are indicated. (C) Fragment ion mass spectrum of the doubly charged precursor of the monophosphorylated peptide NL3(791-800). The inset shows the mass spectrum of the parent phosphopeptide (M_obs_ = 1300.6354, M_calc_ = 1300.6176, relative mass error = 13.7 ppm). Although a contaminating parent peptide ([M+2H^+^]^2+^ = 650.294) was co-isolated for fragmentation with the target parent peptide ([M+2H^+^]^2+^ = 651.325), conclusive N-terminal b-ion (b_2_, b_3_, b_7_ in their non-phosphorylated forms) and C-terminal y-ion (y_5_-y_8_ in their phosphorylated forms) series, together with the neutral loss of the phospho moiety from y_3_, clearly indicates phosphorylation on S799 while excluding the other potential phosphorylation sites of the peptide DYTLTLRRSP. (D) Spot blots assessing the phosphospecificity of the 6808 antibody-peptide mix, targeting the epitope containing phosphorylated S799 on NL3. Phosphorylated and corresponding unphosphorylated peptides spotted on nitrocellulose at increasing amounts (50 ng to 2 μg). Immunoreactivity was measured by infrared fluorescence of dye-conjugated secondary antibody. (E) Western blots of lysates from cerebral cortex (Cx), hippocampus (Hp), striatum (St), thalamus (Th), and cerebellum (Cb), probed for actin (loading control, bottom), NL3 (middle), and phosphorylated at S799 NL3 (NL3-pS, top). Antibody 6808 recognizes a band at the size of NL3 that is absent in *Nlgn3* KO lysates, indicating specific recognition of native phosphorylated NL3. While NL3 was detected in all regions examined, bands corresponding to NL3-pS799 were specifically detected as a major band in hippocampus, and minor bands in cortex and cerebellum.

We sought to identify kinases with substrate specificity for NL3-S799 using *in vitro* assays with purified kinases and substrate peptide (NL3 785-805; RLTALPDYTLTLRRSPDDIPL). We applied this assay in a candidate approach (Extended Data Fig. 3A) and a larger-scale screening approach (Extended Data Fig. 3B) and detected phosphate transfer onto the NL3 substrate peptide with various kinases. However, we did not identify any kinases that displayed substrate-specificity for the S799 residue as compared to other phosphorylatable residues of the peptide. Thus, the kinase responsible for NL3-S799 phosphorylation remains elusive.

**Figure 3.**
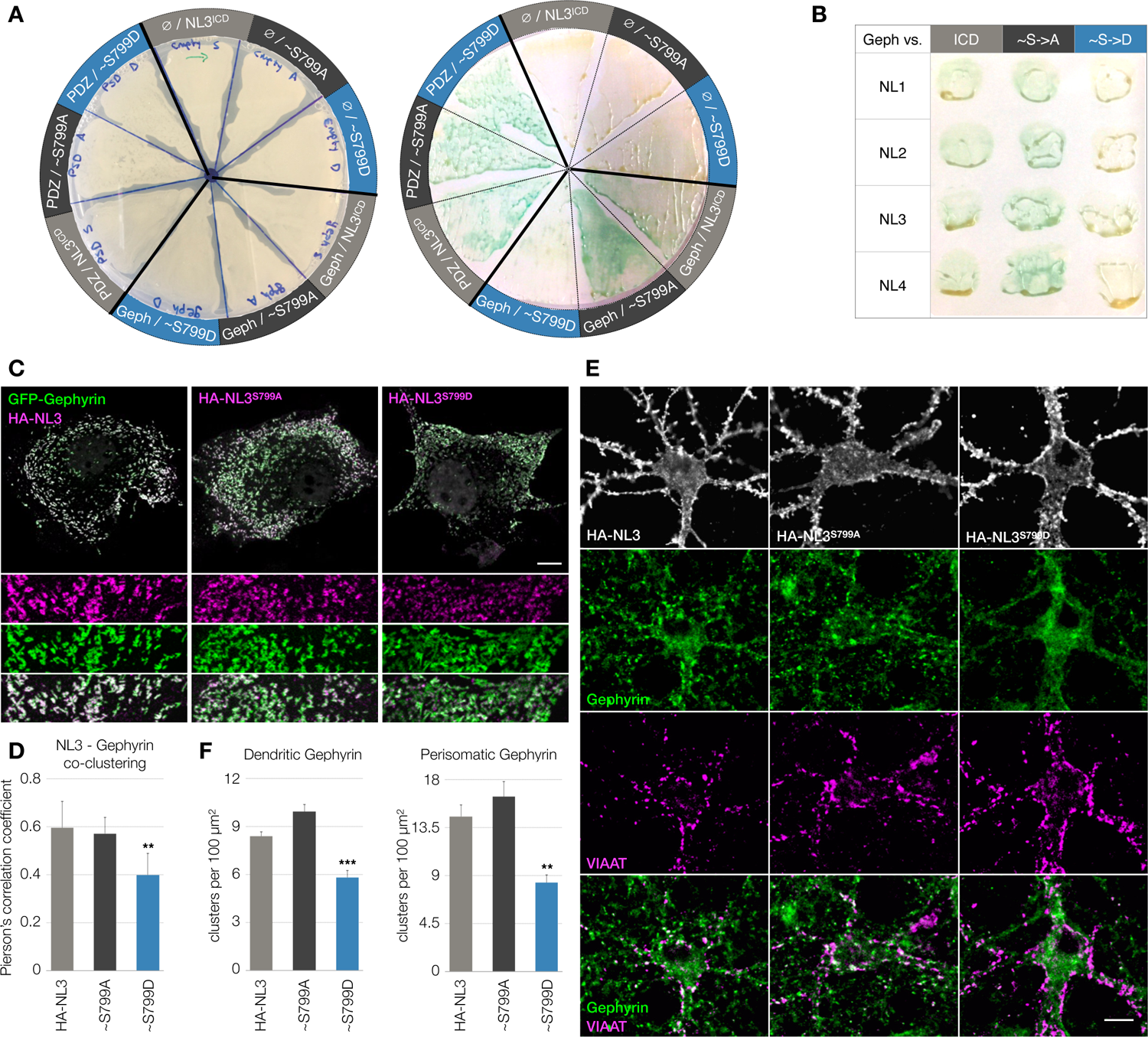
Association of Gephyrin with NLs is affected by phosphomimetic mutation of serine at the Gephyrin-binding site in three exogenous systems. (A) Yeast-two-hybrid assays using NL3 intracellular domains as bait, with WT S799 (NL3^ICD^), phospho-null (∼S799A), or phosphomimetic (∼S799D) variants used against prey constructs of empty vector control (ø), Gephyrin (Geph), or S-SCAM fragment encompassing PDZ domains 1-3 (PDZ). Left: base plate showing comparable growth of transformed yeast cultures across all plate segments. Right: colorimetric β-galactosidase reaction on colony-lift replicate membrane. Positive interaction between NL3 mutants and postsynaptic scaffolds is indicated by cyan color reaction. (B) Yeast-two-hybrid assays with bait intracellular domains from NLs 1-4 in WT (ICD) and corresponding Gephyrin-binding site phospho-null (∼S->A) and phosphomimetic (∼S->D) mutations versus full-length Gephyrin prey constructs. Phosphomimetic mutation at the Gephyrin-binding serine disrupts interaction with Gephyrin. (C) Membrane co-clustering assays in COS7 cells transfected with GFP-Gephyrin (green) and myc-CB2^SH3-^ to induce Gephyrin membrane microaggregates. HA-NL3 (magenta) or its corresponding phospho-null (HA-NL3^S799A^) or phosphomimetic (HA-NL3^S799D^) mutants were transfected to assess co-clustering with Gephyrin membrane microaggregates (scale bar corresponds to 10 μm for main panel and 3.6 μm for inset). (D) Pearson’s correlation coefficients between GFP-Gephyrin microaggregates and surface-stained HA-NL3, HA-NL3^S799A^, or HA-NL3^S799D^; p < 0.01, one-way ANOVA with post-hoc Bonferroni from 3 independent experiments. (E) Expression of HA-NL3, phospho-null (HA-NL3^S799A^), and phosphomimetic (HA-NL3^S799D^) mutants in DIV14 cultured neurons, immunolabeled for endogenous inhibitory postsynaptic marker Gephyrin (green) and inhibitory presynaptic marker VIAAT (magenta), merged in bottom row. Scale bar 10 μm. (F) Quantification of dendritic (p < 0.001, one-way ANOVA with post-hoc Bonferroni, n ≥ 60 cells per condition from 5 independent experiments) and perisomatic (p < 0.01, one-way ANOVA with post-hoc Bonferroni, n ≥ 35 cells per condition from 3 independent experiments) Gephyrin clusters in neurons transfected with HA-NL3 or corresponding mutants. Phosphomimetic mutation at the Gephyrin-binding serine specifically abolishes NL3 interaction with Gephyrin in yeast, hinders NL3-Gephyrin co-clustering in cell lines, and NL3-mediated Gephyrin recruitment in cultured neurons.

We then investigated the endogenous distribution of S799-phosphorylated NL3 (NL3-pS799) and whether it follows region-specific patterns corresponding to NL3 synapse specificities. To do so, we raised phospho-specific antibodies in rabbits immunized against the corresponding NL3-pS799 peptide (RRpSPDDIP). The best performing antibody (6808) displayed moderate preference for the phosphorylated over the non-phosphorylated peptide, which was significantly increased when mixed with an excess of non-phosphorylated peptide (RRSPDDIP) to compete with non-phosphorylated NL3 epitopes. This increased the phospho-specificity of antibody 6808 from approximately 3-fold with antibody alone to 20-fold with the antibody-peptide mix (Fig. 2D), providing us with a critical reagent for the detection of endogenous NL3-pS799.

To investigate the region-specificity of endogenous NL3-pS799, we immunoblotted samples from different brain regions using the phospho-specific 6808 antibody-peptide mix. We validated detection specificity and determined whether homologous phosphorylation occurs in other NLs by blotting samples from WT as well as NL3 KO mice as controls. We blotted samples from five brain regions (neocortex, hippocampus, striatum, thalamus, and cerebellum) from both WT and KO genotypes on the same membranes. Actin across all lanes indicated equal total protein load. NL3 immunoreactivity appeared as a specific band with electrophoretic mobility corresponding to 100-110 kDa, which was absent in NL3 KO samples. Similar levels of total NL3 immunoreactivity were detected across WT lanes from all brain areas examined (Fig. 2E).

Having validated the preparation, we immunoblotted with the 6808 antibody-peptide mix and identified specific NL3-pS799 immunoreactive bands in the anticipated 100-110 kDa range, displaying robust intensity in WT hippocampus, and moderate intensity in WT cortex and cerebellum. Equivalent bands were absent from corresponding NL3 KO samples, confirming that the phospho-specific signal comes from NL3. Given that the Gephyrin-binding motif is highly conserved among NLs, the complete loss of phospho-specific bands in NL3 KOs shows that NL1 and NL4 are not phosphorylated at the homologous site, indicating that this phosphorylation mechanism is specific to NL3.

Importantly, while NL3 was robustly detected in all areas tested, phospho-specific NL3-pS799 bands seen in hippocampus, neocortex, and cerebellum were not seen in striatum or thalamus (Fig. 2E), the latter being areas in which NL3 puncta colocalize with Gephyrin at inhibitory synapses (Fig. 1). Indeed, the region specificity of the NL3-pS799 band corresponds to the pattern of NL3 localization at excitatory synapses, with NL3 being present at the majority of excitatory synapses in hippocampus, and in more limited subsets of excitatory synapses in neocortex and cerebellum. From these results we conclude that the newly identified NL3-pS799 represents an endogenous phosphorylation variant that quantitatively correlates with the region-specific fractions of NL3 that localize to excitatory synapses.

### Phosphomimetic mutation of S799 in the Gephyrin-binding site of NL3 inhibits association with Gephyrin

S799 is located within the R797-T812 epitope that corresponds to the minimal Gephyrin-binding site^19^, indicating that phosphorylation at this site may regulate NL3-Gephyrin interaction. We detected NL3-pS799 specifically in areas where NL3 does not colocalize with Gephyrin, while NL3 is unphosphorylated in subcortical areas where NL3 and Gephyrin do colocalize. We thus hypothesized that S799 phosphorylation of NL3 inhibits its association with Gephyrin. To test this, we produced phosphomimetic S799D (NL3^S799D^) and phospho-null S799A (NL3^S799A^) mutants to genetically mimic phosphorylated and unphosphorylated variants, and used these in a series of experiments to assess NL3 association with Gephyrin in yeast, cell lines, cultured neurons, and in the brain.

We first used yeast-two-hybrid to assay NL3^S799D^ and NL3^S799A^ cytoplasmic domain interactions with Gephyrin or with PDZ domains of S-SCAM, a scaffold protein of excitatory postsynapses. Consistent with our hypothesis, binding of NL3 to Gephyrin was abolished by phosphomimetic NL3^S799D^, while PDZ-mediated interactions were left intact (Fig. 3A). Phospho-null NL3^S799A^ interacted with both Gephyrin and PDZ domains (Fig. 3A). Corresponding results were obtained with homologous mutants of all other NL cytoplasmic domains (Fig. 3B, Extended Data Fig. 4). These data indicate that serine phosphorylation of the Gephyrin-binding site of NLs prevents the interaction with Gephyrin.

We next examined the association of full-length NL3 and its NL3^S799D^ and NL3^S799A^ mutants with Gephyrin membrane clusters in heterologous mammalian cells. Such clustering assays have been used extensively to examine the association of Gephyrin with membrane proteins, including glycine receptors^37^, GABA_A_ receptors, and NLs^19^. HA-tagged NL3 constructs were expressed in COS7 cells together with constructs expressing GFP-Gephyrin and myc-tagged Collybistin lacking its autoinhibitory SH3 domain (myc-CB2^SH3-^), a constitutively active form of Collybistin that induces Gephyrin membrane clustering in heterologous cells^19, 38, 39^. GFP-Gephyrin membrane clusters induced pronounced co-clustering of HA-NL3 (Fig. 3C-4D), which we quantified using cross-correlation analysis as done previously^19^. Phosphomimetic HA-NL3^S799D^ displayed significant reduction in co-clustering with Gephyrin by approximately one-third compared to WT or HA-NL3^S799A^ (Fig. 3D), consistent with disruption of the Gephyrin-binding site by S799 phosphorylation.

We went on to test the effects of these constructs in cultured neurons. Overexpression of WT HA-NL3, as well as phospho-null HA-NL3^S799A^ increased the number of Gephyrin clusters compared to phosphomimetic HA-NL3^S799D^ (Fig. 3E-3F). This abating effect of the phosphomimetic mutant on Gephyrin clustering is consistent with a loss of its ability to interact with Gephyrin at the neuronal plasma membrane. Together, our results obtained with cultured cells support the notion that phosphorylation of S799 in the Gephyrin-binding site of NL3 perturbs its association with the Gephyrin scaffold at inhibitory postsynapses.

### Phosphomimetic mutation of S799 localizes NL3 to excitatory synapses *in vivo*

We finally investigated whether S799 phosphorylation determines synaptic localization of NL3 *in vivo*. We generated constructs expressing HA-tagged NL3 variants under a synapsin promoter and delivered these by *in utero* electroporation to the mouse cortex. We co-electroporated constructs expressing fusion proteins that label excitatory and inhibitory postsynaptic scaffolds to determine subcellular localization of HA-NL3 variants. Analysis was performed in three-week old animals to localize HA-NL3 WT, S799A phospho-null, and S799D phosphomimetic variants in relation to excitatory postsynapse marker PSD95-RFP^40^ and the inhibitory postsynapse nanobody Gephyrin.FingR-GFP^41^ (Extended Data Fig. 6A). Cross-correlation analysis of the HA signal with RFP and GFP signals was used to ratiometrically quantify excitatory vs. inhibitory synapse localization of each HA-tagged NL3 variant.

HA-NL3 localization in cortex strongly correlated with PSD95 marker at excitatory postsynapses over Gephyrin marker at inhibitory postsynapses (Extended Data Fig. 6A), consistent with our findings of endogenous NL3 excitatory localization in cortex (Fig. 1). Similarly, the phosphomimetic S799D variant showed strong specificity for excitatory postsynapses, in line with our findings that NL3 in cortex is phosphorylated at S799 (Fig. 2). The phospho-null S799A variant showed marked reduction in preference for excitatory postsynapses, but it did not show specificity for inhibitory over excitatory postsynapses, as we had hypothesized (Extened Data Fig. 6B). This reduced specificity for excitatory postsynapses (E-I of 0.2 ± 0.039 for S799A compared to 0.47 ± 0.057 and 0.46 ± 0.023 for WT and S799D, respectively) may reflect an increased association of HA-NL3^S799A^ with Gephyrin by preventing endogenous phosphorylation at the S799 position. However, this effect was partial and not sufficient to confer specificity for inhibitory synapses.

We and others showed previously that NLs exist as dimers on the cell surface^32, 42–46^. We thus posited that exogenous HA-tagged constructs, including the phospho-null HA-NL3^S799A^, may dimerize with endogenous NL3 that is phosphorylated in cortex, resulting in mixed HA-NL3^S799A^-NL3^WT^ dimers that produce the observed mixed E-I specificity of HA in the *in vivo* experiments (Extended Data Fig. 6). To circumvent confounding dimerization with endogenous NL3, we developed CRISPR reagents targeting the *Nlgn3* gene for somatic cell knockout in electroporrated neurrons. We tested Cas9 with candidate guide RNAs (gRNAs) and selected the target sequence with the highest efficiency using an *in vitro* cleavage assay^47^. We confirmed efficacy of Cas9 with gRNA::*Nlgn3* for producing frame-shift insertions and deletions (indels) in the *Nlgn3* locus of mouse Neuro-2A cells (Fig. 4).

**Figure 4.**
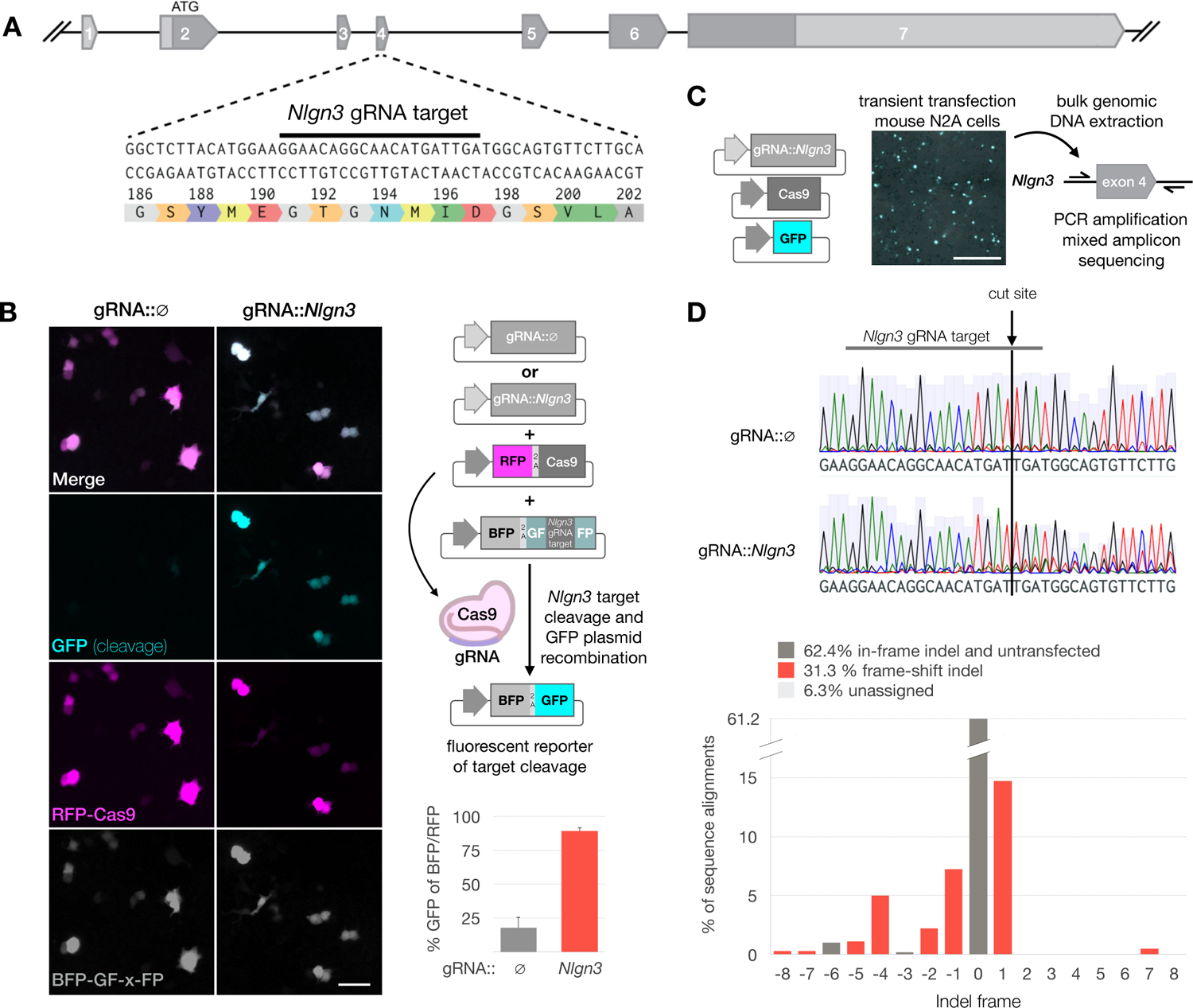
Construction and validation of CRISPR knockout plasmids for NL3. (A) Structure of the *Nlgn3* gene. Exons are numbered with annotation of ATG and open reading frame in dark grey. Closeup of *Nlgn3* nucleotide sequence and translation of the exon 4 region targeted by *Nlgn3* gRNA indicated by black bar. (B) Split GFP assay to visualize cleavage of the *Nlgn3* gRNA target sequence. Neuro-2A cells imaged 48h post transfection with three plasmids as schematized: i) gRNA plasmid containing either control (ø) or gRNA targeting *Nlgn3*, ii) plasmid expressing Cas9 and RFP (magenta), and iii) cleavage reporter plasmid expressing BFP (grey) and containing GFP sequence split by *Nlgn3* gRNA target sequence. Successful cleavage of the *Nlgn3* gRNA target sequence by Cas9+gRNA results in recombination of the reporter plasmid producing GFP fluorescence (cyan). Scale bar 10 μm. Graph shows the percentage of BFP and RFP cells that show cleavage reporter GFP expression in control gRNA 17.9% (± 7.7% S.E.M.) and *Nlgn3* gRNA 89.3% (± 2.5% S.E.M.) transfections. Averages from N=3 biological replicates, each averaged from n=9 imaged fields. Scale bar 100 μm. (C) Genomic DNA sequencing of *Nlgn3* exon 4 locus to assess and confirm deleterious insertion-deletion (indel) production. Mouse Neuro-2A cells co-transfected with Cas9 plasmid, GFP reporter plasmid (cyan), and either control or *Nlgn3* gRNA plasmid assayed for cleavage in B. Scale bar 100 μm. Bulk genomic DNA was isolated 7 days post-transfection and used as template for PCR of *Nlgn3* exon 4. (D) Chromatographs of mixed amplicon Sanger sequencing from control gRNA::ø (top) or gRNA::*Nlgn3* (bottom) transfected Neuro-2A cells from C. Control chromatographs were used as reference for unmixing of indel fitting of the mixed signal in the gRNA::*Nlgn3* sample. Histogram shows assignment of indel frame frequency. Bins of deleterious frame-shift indels labeled in red, together make up 31.3 % of total reads. In-frame indels or wild type sequence bins are marked in grey. They comprised 62.4% of reads, which is close to the range anticipated for untransfected cells.

Having validated NL3 KO efficacy of the CRISPR plasmids, we included them in subsequent iterations of the *in utero* electroporation experiments to assess the synaptic localization of HA-NL3 variants on a NL3 KO background in mouse cortex (Fig. 5A). To do so, we modified the HA-NL3 constructs with silent mutations at the site targeted by the gRNA::*Nlgn3* to confer CRISPR-Cas9 resistance and enable replacement of endogenous NL3 with the exogenous variants. Indeed, phospho-null HA-NL3 displayed robust inhibitory synapse specificity (E-I −0.27 ± 0.066) in the absence of endogenous NL3, while HA-NL3 WT (E-I 0.31 ± 0.053) and phosphomimetic S799D (E-I 0.47 ± 0.045) variants retained their excitatory synapse specificity (Fig. 5B-C). These results show that S799 phosphomimetics at the Gephyrin-binding site of NL3 determine the scaffold specificity and synaptic localization of NL3 *in vivo* and support the notion that NL3 localizes to excitatory synapses in cortex due to S799 phosphorylation.

**Figure 5.**
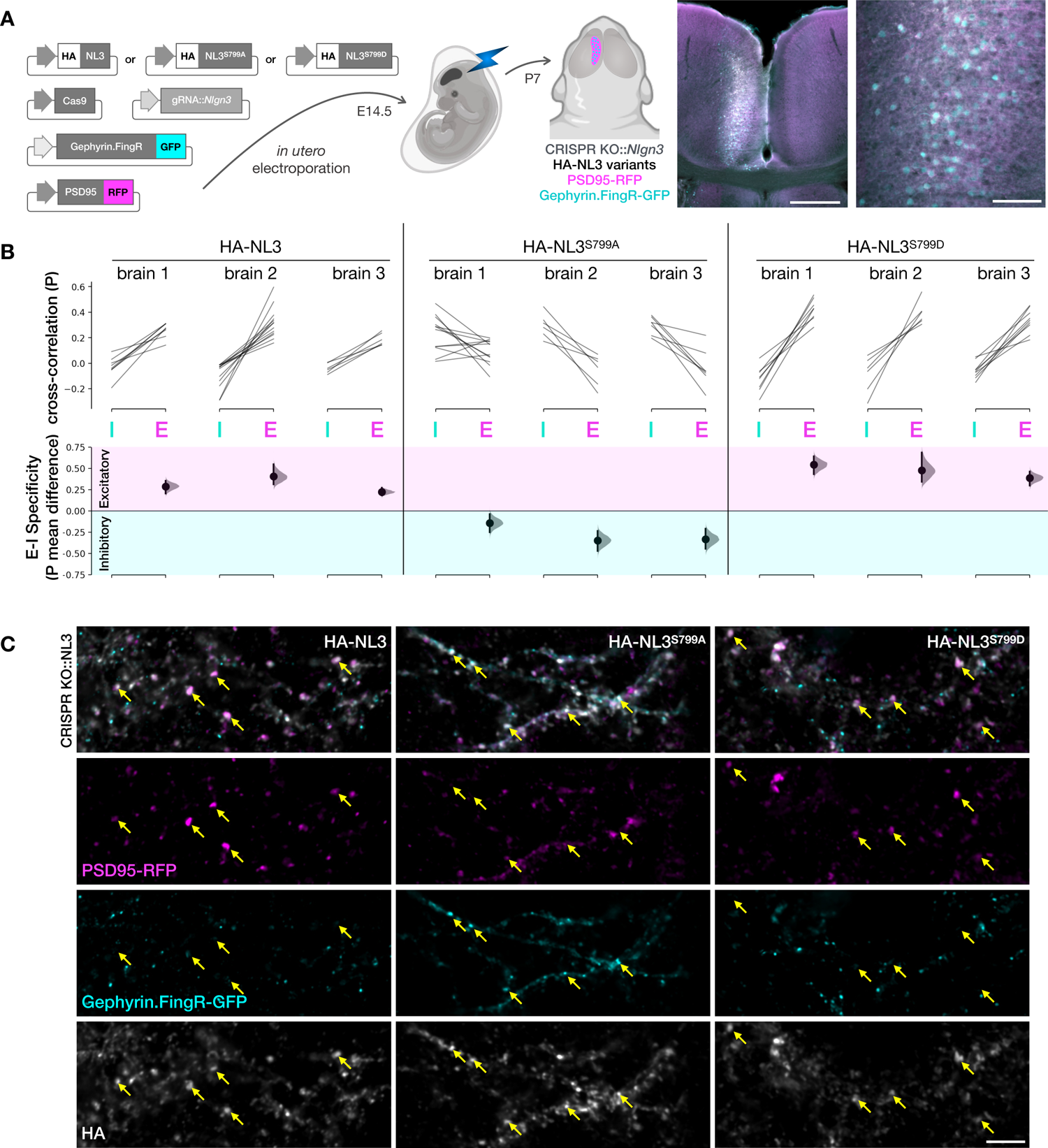
Phosphomimetic mutation of the Gephyrin-binding site on NL3 determines excitatory versus inhibitory synapse specificity in vivo. (A) Schematic of in utero electroporation approach to determine synapse localization of tagged NL3 variants in a knockout background mouse cortex. Plasmid cocktails were used to test each HA-tagged NL3 variant (WT, S799A phospho-null, and S799D phosphomimetic) consisting of Cas9 and gRNA::*Nlgn3* to knock out endogenous NL3, Gephyrin.FingR-GFP (cyan) to label inhibitory postsynapses, and PSD95-RFP (magenta) to label excitatory postsynapses. Plasmid cocktails were electroporated into E14.5 mouse embryos and brains imaged at P7. Example images are shown of electroporation site in two magnifications. Scale bars 500 μm (left), and 100 μm (right). (B) Specificity of excitatory versus Inhibitory postsynapse localization was quantified in three brains for each HA-NL3 variant on a *Nlgn3* CRISPR KO background, as well on a WT background (see Extended Data Fig. 6). Localization was quantified based on cross-correlation analysis of the HA signal to GFP inhibitory (I) and RFP excitatory (E) synapse marker signals. Top plot shows average cross-correlation values (P) in paired E-I analyses from each brain section in each of three brains per variant. Bottom plot shows paired mean differences of excitatory versus inhibitory correlations per brain in Cumming estimation plots. Effect size is shown with sampling distributions and vertical bars indicating 95% confidence interval. Positive values indicate specificity of HA-NL3 variants for excitatory postsynapses (magenta), while negative values indicate specificity for inhibitory postsynapses (cyan). In the absence of endogenous NL3, cortically expressed HA-NL3 showed excitatory synapse specificity for WT (0.31 ± 0.053) and phosphomimetic S799D mutant (0.47 ± 0.045) variants, while phospho-null S799A mutant HA-NL3 showed inhibitory synapse specificity (−0.27 ± 0.066). Values are E-I mean difference from n=6 to 13 sections (depending on electroporation field size) from each brain, with deviations in SEM for N=3 brains per group. (C) Representative images in electroporation fields of cortex after *Nlgn3* knockout (CRISPR KO::NL3) showing merged (top) and individual localization of HA-NL3 variants in white, inhibitory postsynapse marker Gephyrin.FingR-GFP in cyan, and excitatory postsynapse marker PSD95-RFP in magenta. Yellow arrows indicate bona fide synaptic puncta with HA colocalizing with excitatory (WT and S799D columns) or inhibitory (S799A column) postsynapse markers. Scale bar 5 μm. WT and phosphomimetic S799D mutant HA-NL3 localizes to excitatory synapses in cortex, while phospho-null S799A mutant HA-NL3 inappropriately localizes to inhibitory synapses in cortex.

## DISCUSSION

NL3 is the most abundant NL in the rodent brain^34^ and currently the focus of intense research owing to its critical functions in synapse development, maturation, and transmission^3–5, 48^, as well as its emerging roles in autism^1, 2,49^and cancer^7, 8^. In this work we investigated the subcellular specificities of NL3 and the cell biological mechanisms that determine them in their native neuronal contexts.

Synapse-type specificities are well documented for NL1, NL2, and NL4 in rodents^14, 17, 18, 22, 23, 34^. The data presented here demonstrate that NL3 has regional synapse-type specificities: excitatory synapses in the cerebral cortex, hippocampus, and areas of cerebellum; and inhibitory synapses in olfactory bulb, basal ganglia, thalamus, brain stem, and the granule cell layer of cerebellum. Focused investigations of NL3 localization have reported it at inhibitory synapses in striatum^21^ and retina^26^, and in subsets of excitatory and inhibitory synapses in hippocampus^50, 51^. These findings illustrate a versatile and highly regulated role of NL3 at different synapse subtypes, and highlights the need for future systematic study at each of these regions.

### Mechanisms that determine scaffold specificities of NLs

All NLs contain binding sites for the key scaffold proteins at inhibitory^19^ and excitatory postsynapses^14, 35^ (Fig. 2 and Fig. 6). In heterologous systems, NLs often lose native specificities and associate non-selectively with both postsynaptic scaffold types (e.g. Fig. 3A and Extended Data Fig. 1). *In vivo*, however, NLs display high synapse-type specificities, indicating endogenous mechanisms regulating affinity of a given NL to either excitatory or inhibitory postsynaptic scaffolds. Interestingly, the mechanisms regulating synapse-type specificity appear to be distinct for different NLs.

The mechanism by which NL2 and mouse NL4 associate with Gephyrin involves their ability to activate Collybistin, a Gephyrin accessory protein, which NL1 and NL3 do not^19, 39^. The specificity of NL1 for excitatory synapses results from constitutive phosphorylation of a tyrosine residue that blocks Gephyrin binding^36, 52^. Here, we identify a new phosphorylation-based mechanism that determines the synapse-type specificity of NL3, which –unlike mechanisms for the other NLs– displays striking region-specificity.

Using mass spectrometry on brain lysates, we detected a previously unknown phosphorylated serine residue in the Gephyrin-binding site of NL3. We determined with phosphomimetic mutants that the corresponding phosphoserine prevents association of NL3 with Gephyrin, similar in function to the phosphotyrosine in NL1. Generation and validation of phospho-specific antibodies against the phospho-serine epitope confirmed that phosphorylation occurs in areas where NL3 localizes to excitatory synapses. Using an *in utero* NL3 replacement strategy, we determined that phosphomimetic NL3 localized to excitatory synapses in cortex, and phospho-null NL3 inappropriately localized to inhibitory synapses in cortex (Fig. 5), supporting that phosphorylation of the Gephyrin-binding site is necessary and sufficient for NL3 localization to excitatory synapses *in vivo*.

The Gephyrin-binding motif is highly conserved across all NLs, and the Gephyrin-binding serine residue is present in NL1, NL3, and NL4. Interestingly, NL2 has an alanine in the corresponding position (Fig. 2A), effectively making it a phospho-null variant that hence always binds Gephyrin, in agreement with NL2 being found exclusively at inhibitory synapses^18, 19^. The peptide we used to produce the phospho-specific antibody equivalently represents conserved epitopes in NL1 and NL4. However, the phospho-specific band we see in brain lysates of WT animals is absent in NL3 KO lysates, demonstrating that only NL3 is phosphorylated at that site at detectable levels (Fig. 2E). These results add to an emerging theme of NL isoform-specific phosphorylation events at distinct sites, including tyrosine and threonine phosphorylation of NL1^36, 53, 54^, serine phosphorylation of NL2^55^, and threonine phosphorylation of NL4^56^. Together, these data indicate that distinct mechanisms have evolved to regulate the synapse-type specificities of the four NLs, which in the case of NL3 is region-specific.

Importantly, NLs appear on the neuronal surface as dimers^32, 42–46^. Our experiments are in support of *in* vivo dimer formation, as exogenous NL3 variants showed less pronounced mutant-specific effects on a WT background (Extended Data Fig. 6) than on a NL3 KO background (Fig. 5). NL3 forms both homodimers and NL1-NL3 heterodimers in culture (Poulopoulos 2012). While no NL2-NL3 dimers were detected, complexes of NL2 and NL3 were co-immunoprecipitated from brain lysates^25^, indicating higher order postsynaptic complexes containing both NL2 and NL3. Considering our findings together with findings of NL1 tyrosine phosphorylation^36, 54^, a model emerges where phosphorylated NL dimers populate excitatory postsynapses, and unphosphorylated NL dimers populate inhibitory postsynapses (Fig. 6). These three features together, namely (i) intrinsic specificities, (ii) distinct dimer combinations, and (iii) post-translational modifications, including Gephyrin-biding site phosphorylation, contribute to an enriched pallet of NL diversity at synapse subtypes.

**Figure 6.**
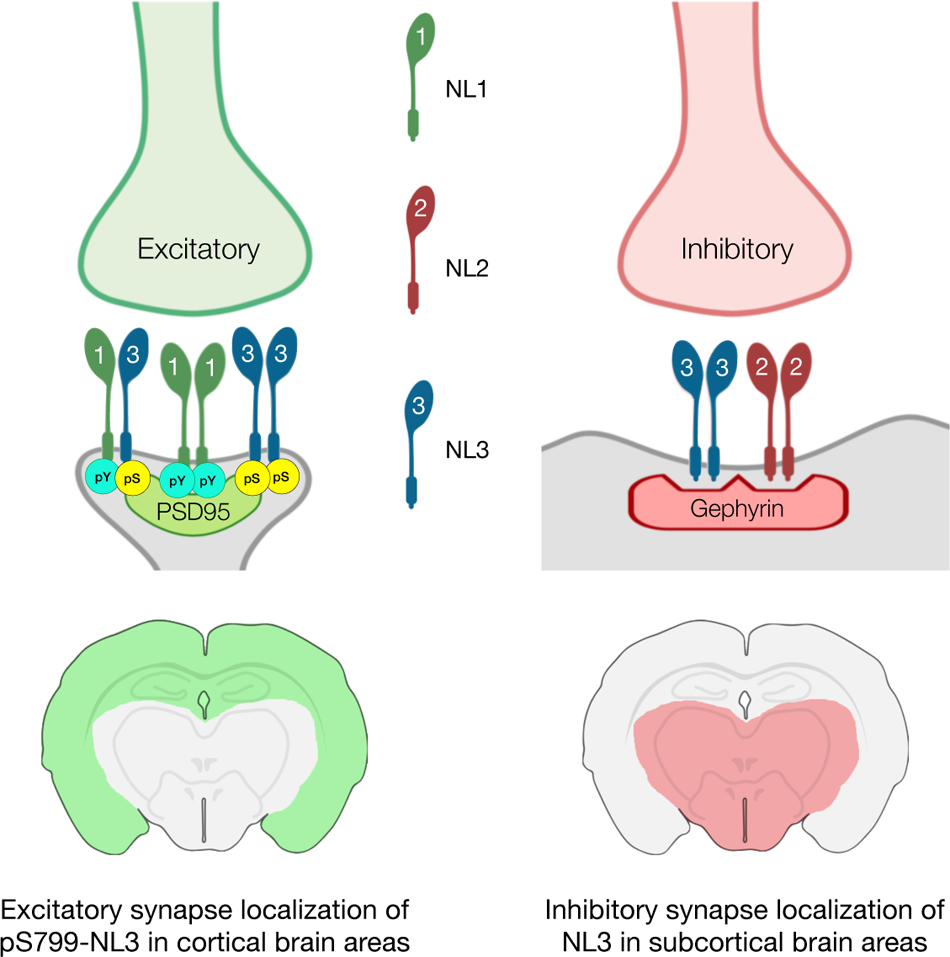
Model of NL3 region-specific, phosphorylation-dependent synapse localization. Schema of the proposed model of NL postsynaptic localization synthesizing the current NL3 findings with previous findings^14, 19, 25, 35, 45^. In cortical regions of the brain (shaded green in coronal brain section schema), including the cerebral cortex and hippocampus, NL3 is phosphorylated at its Gephyrin-binding site leading to association with PSD95 and localization to excitatory postsynapses. In subcortical regions of the brain (shaded red in schema), including basal ganglia, thalamus, and brainstem, NL3 remains unphosphorylated at its Gephyrin-binding site leading to association with Gephyrin and localization to inhibitory postsynapses. Synapse-type specificities of NL3 are schematized together with other NLs in dimer states as previously described. Gephyrin-binding sites are phosphorylated in NL dimers comprised of NL3-pS799 (pS) and NL1-pY782 (pY) at excitatory postsynapses. NL3 dimers are S799 unphosphorylated together with NL2 dimers at inhibitory synapses.

### NL3 specificities in mouse models of disease

Studies in NL3 mutant mice have identified strikingly diverse synaptic phenotypes, indicating diverse function of NL3 in different parts of the brain. Phenotypes are largely consistent with our localization data, though we did not investigate synapse subtypes at the circuit level. In hippocampus, where we find NL3 heavily phosphorylated (Fig. 2E) and at excitatory synapses (Fig. 1B-C), NL3 KO caused excitatory synapse phenotypes in CA1^57, 58^, dentate gyrus^59^, and stratum oriens^60^, consistent with our findings. Other studies focusing on subtype-specific phenotypes^11^ and cell type-specific localization of NL3 in hippocampus^50, 51^ indicate more refined specificities, including at inhibitory CCK synapses, which require focused investigations into putative roles of phosphorylated NL3 at those subtypes. In the basal ganglia, where we find unphosphorylated NL3 to localize to inhibitory synapses (Fig. 2D and 3E), recordings from D1 medium spiny neurons of ventral striatum show loss of inhibitory postsynaptic currents in NL3 KOs^10^, consistent with our findings. On aggregate, region-specific NL3 specificities will be key to interpreting synaptic effects in NL3 mouse lines, including autism models^12^.

NL3 is unusual among synaptic adhesion molecules because it switches scaffold specificities and shifts localization between excitatory and inhibitory synapses. This is particularly intriguing in light of the proposed role of excitation/inhibition imbalance in autism^61–63^. While monogenic forms of autism are rare, polygenic and other etiologies may converge on the same elements of brain connectivity where coordinated development of excitatory and inhibitory synapses is crucial.

The involvement of NL3 in the tumor microenvironment of the brain is surprising both in mechanism and potency. NL3 is shed by activity at the synapses it localizes to and promotes the spread of glioma in mice^7^. Conversely, xenografted patient-derived tumor cells fail to grow in NL3 KO mice, with no other NL showing such effects^8^. 86% of glioma cases are in cortex^64^ where we find NL3 to be excitatory, indicating that regional specificities of NL3 may affect regional glioma spread. Pharmacologically targeting activity at synapse types that contain NL3 based on glioma location may provide a strategy to contain glioma growth. Additionally, CRISPR agents disrupting NL3 expression, such as those we report here, can be developed into putative therapeutic agents to impede relapse after tumor resection.

These avenues of future research, along with refined determination of NL3 synapse subtype specificities, will help understand how atypical connectivity during brain development causes autism, and may provide new cancer treatment options.

## ACKNOWLEDGEMENTS

This study was supported by the German Research Foundation (SFB1286/A09, N.B.), Germany’s Excellence Strategy (EXC 2067/1-390729940, N.B.), the Ministero dell’Istruzione, dell’Università, e della Ricerca (MUR Project Dipartimenti di Eccellenza 2018-2022, Department of Neuroscience “Rita Levi Montalcini”; An.P., M.S.-P.), and the Office of the Director of the National Institutes of Health (award number DP2MH122398; Al.P.). L.P.T. was supported by a Marie Curie Fellowship (grant #274972). We are grateful to H. Betz (Heidelberg), J. Miyazaki (Osaka), S. Jamain (Göttingen, Paris), A.M. Craig (Vancouver), T. Dobbie (Vancouver), and D. Arnold (Los Angeles) for generously providing plasmids. We thank S. Wenger, K. Hellmann, and L. van Werven (Göttingen) for technical support.

## AUTHOR CONTRIBUTIONS

Conceptualization, N.B., L.P.T., and Al.P.; Methodology, L.P.T., K.D., H.U., O.J., N.B., Al.P. Investigation, L.P.T., B.A., An.P., K.D., T.S., A.J.R., C.D.R., S.N.K., G.W.B., M.C.A., O.Y., D.K., M.H., H.H., P.R.L., L.D., M.S.-P., J.J.E.C., Al.P.; Formal Analysis L.P.T., B.A., An.P., K.D., T.S., A.J.R., C.D.R., S.N.K., G.W.B., H.H.H., Al.P.; Writing – Original Draft, L.P.T., Al.P.; Writing – Review & Editing, all authors; Visualization, L.P.T., B.A., An.P., A.J.R., K.D., A.P.; Supervision L.P.T., M.S.-P., H.U., O.J., N.B., Al.P.; Project Administration N.B. and Al.P.; Funding Acquisition M.S.-P., N.B, and Al.P.

## INCLUSION AND DIVERSITY STATEMENT

One or more of the authors of this paper self-identifies as an underrepresented ethnic minority in science. One or more of the authors of this paper self-identifies as a member of the LGBTQ+ community. One or more of the authors of this paper received support from a program designed to increase minority representation in science.

---

**Extended Data Figure 1.**
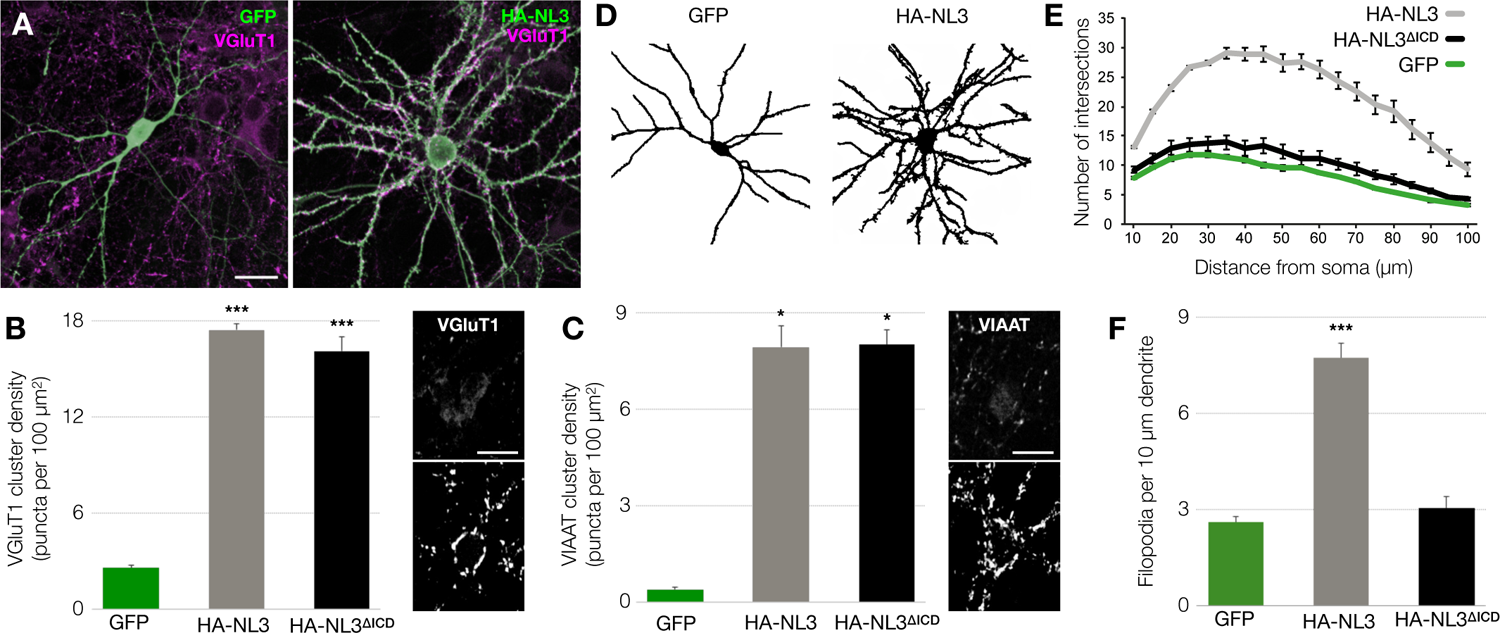
Non-specific dendritogenic and synaptogenic effects of NL3 overexpression in cultured neurons. (A) Cortical neurons at DIV7 overexpressing control GFP or HA-NL3 (green), immunolabeled for the excitatory presynaptic marker VGluT1 (magenta); scale bar, 20 μm. (B-C) Quantification of excitatory VGluT1 (B) and inhibitory VIAAT (C) punctum density onto neurons transfected with control GFP, full-length HA-NL3, and HA-NL3/ΔICD. Sample images of neurons overexpressing HA-NL3 (bottom inset) or control GFP (top inset); scale bar, 20 μm; ANOVA p< 0.05 (*) and p < 0.001 (***) indicated, n ≥ 36 cells per condition from 3 independent experiments. (D) Representative tracing of neurite arborization of neurons transfected with control GFP or HA-NL3. (E) Quantification of dendritic arborization by plotting the number of neurites intersecting concentric circles at 10 μm increments from the soma (Sholl analysis). (F) Density of filopidia per 10 μm of dendrite length from neurons transfected with control GFP, full-length HA-NL3, or cytoplasmic truncation HA-NL3/ΔICD; ANOVA p < 0.001 (***) indicated, n ≥ 60 cells per condition from 4 independent experiments. Exogenous NL3 overexpression promotes dendritic complexity via its cytoplasmic domain and promotes non-specific excitatory and inhibitory synaptogenesis via its extracellular domain.

**Extended Data Figure 2.**
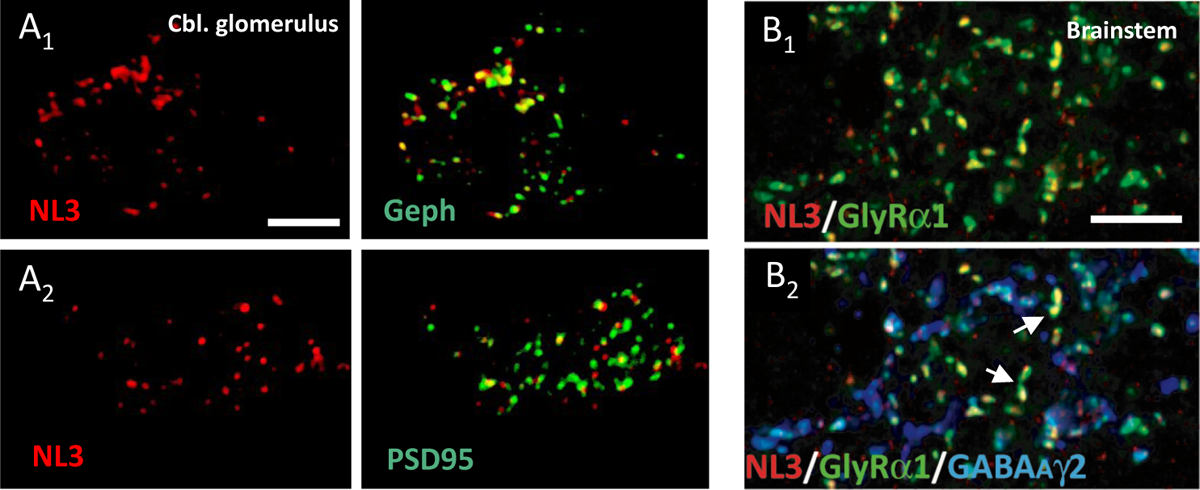
NL3 synaptic localization in cerebellar glomeruli and brainstem. (A) NL3 localizes to both glutamatergic and GABAergic synapses in cerebellar glomeruli. Confocal images showing colocalization of NL3 with both PSD95 (A_1_) and Gephyrin (A_2_) at glomerular synapses. Scale bar 3 μm. (B) Triple-labeling for NL3 (red), GlyR1 (green) and GABA_A_Rγ2 (blue) in the brainstem. Note that NL3 is present at synapses containing both glycine and GABA_A_ receptors. In some cases, NL3 was associated with purely glycinergic synapses lacking GABA receptors (arrows). Scale bar 7 μm.

**Extended Data Figure 3.**
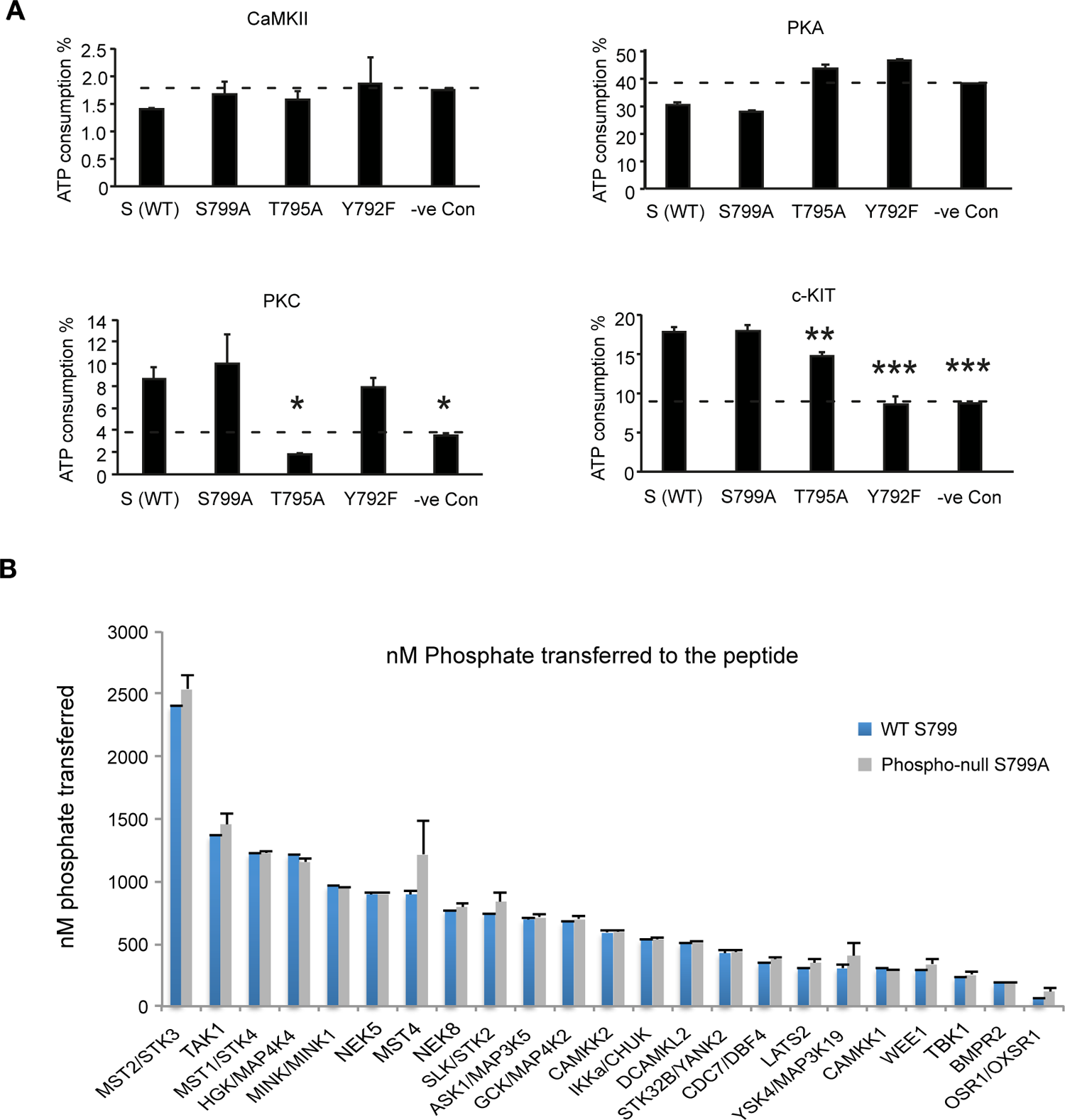
*In vitro* kinase assays do not identify selective NL3 S799 kinase. (A) *In vitro* colorimetric kinase activity assays testing a selection of purified kinases (CaMKII, PKA, PKC, and c-kit) on a 21-residue substrate peptide corresponding to NL3 785-805 (RLTALPDYTLTLRRSPDDIPL) and control peptides with corresponding alanine mutations of putative phosphorylatable positions S799, T795, and Y792. Baseline signal indicated as “-ve control”. Results are means of three data sets. No specific phosphorylation of position S799 was detected. (B) High-throughput *in vitro* radiolabeled phosphate kinase assays measuring phosphate transfer onto the NL3 785-805 peptide and corresponding S799A negative control against 365 individual kinases. The graph shows top kinases ranked in order of phosphate transferred. Experiment was carried out in single dose duplicates. None of the kinases displayed specific phosphorylation of S799.

**Extended Data Figure 4.**
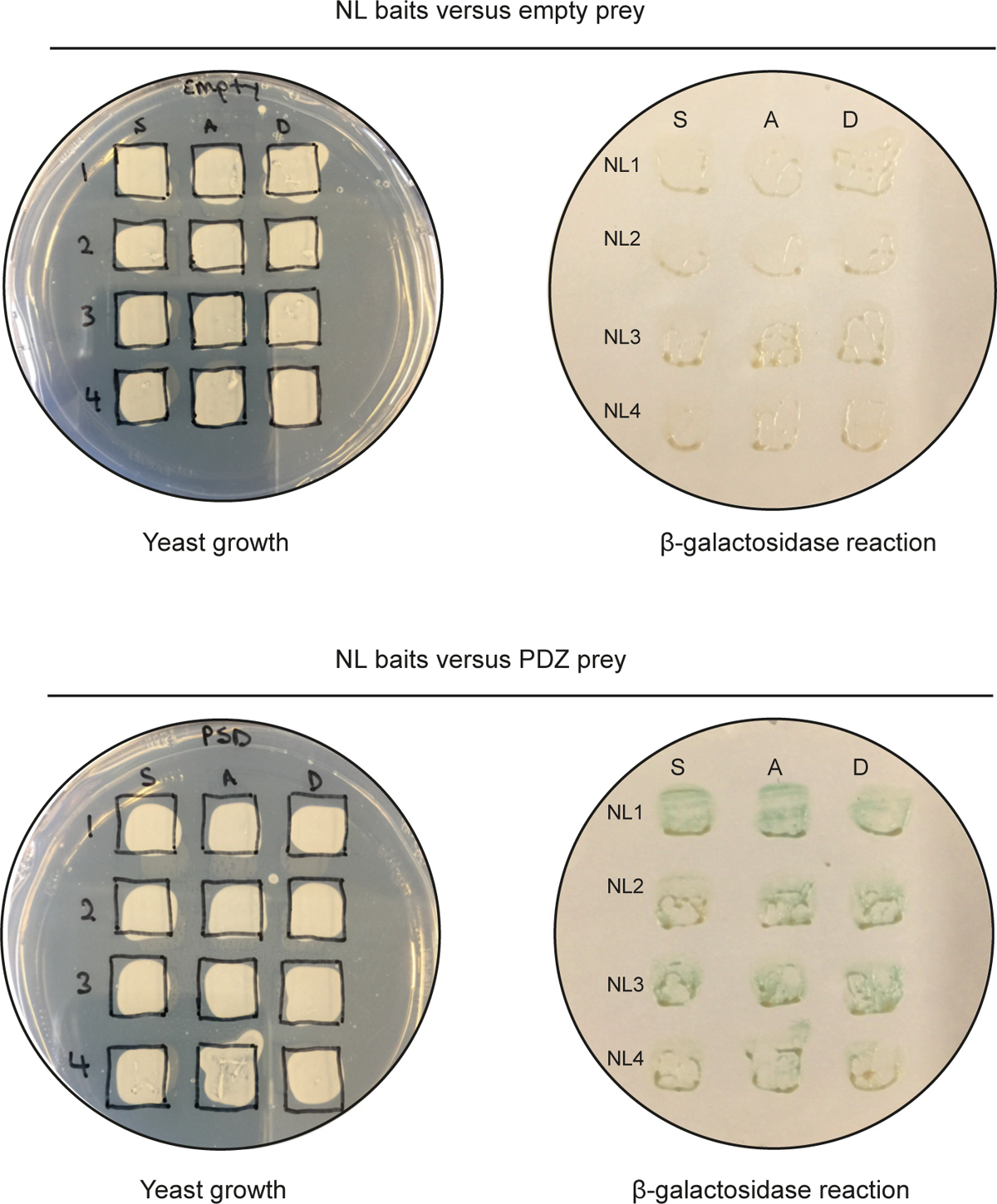
Yeast-two-hybrid of NL intracellular domains versus PDZ domains. Yeast-two-hybrid culture plates (left) and β-galactosidase filter assays (right) of NL1-4 intracellular domain bait constructs in their WT forms (S) and their respective phosphomimetic (D) and phospho-null (A) mutants of the Gephyrin-binding site serine residue (S799 in NL3) versus empty control prey construct (upper) and prey construct with an S-SCAM fragment encompassing PDZ domains 1-3 (PDZ, bottom row). PDZ domain interactions remain unaffected by serine mutation in all NLs.

**Extended Data Figure 5.**
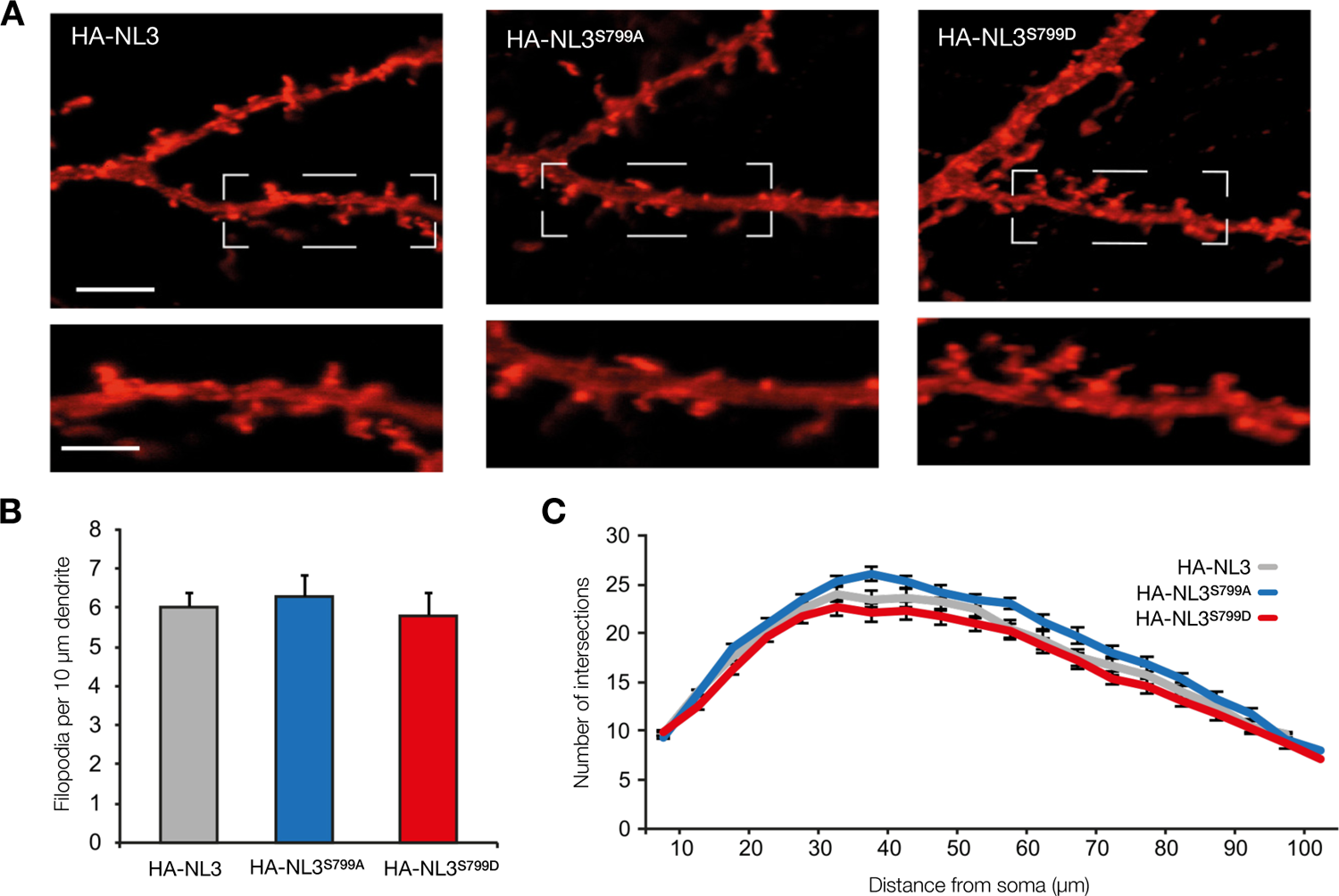
S799 variants of NL3 do not affect dendritic filopodia or arborization. (A) Representative images of dendrites in cortical neurons transfected with HA-NL3 variants of the S799 residue, showing filopodia quantified in (B). Scale bar 5 μm, Areas highlighted in dashed white boxes are shown as insets below. Scale bar 5 μm, insets 3 μm. (B) Quantification of the density of filopodia in neurons transfected with HA-NL3 WT, (grey) phospho-null S799A (blue), and phosphomimetic S799D (red) variants. At least three dendritic segments were analyzed per cell, n ≥ 33 cells per condition from N=3 independent experiments. ANOVA p > 0.05 in all cases. (C) Quantification of dendritic arborization in neurons transfected with HA-NL3 WT, (grey) phospho-null S799A (blue), and phosphomimetic S799D (red) variants. The graph plots the number of neurites intersecting concentric circles at 10 μm increments from the soma (Sholl analysis). n ≥ 90 cells per condition from N=6 independent experiments. Error bars are SEM. No effect of S799 mutation was seen on dendritic filopodia or arborization.

**Extended Data Figure 6.**
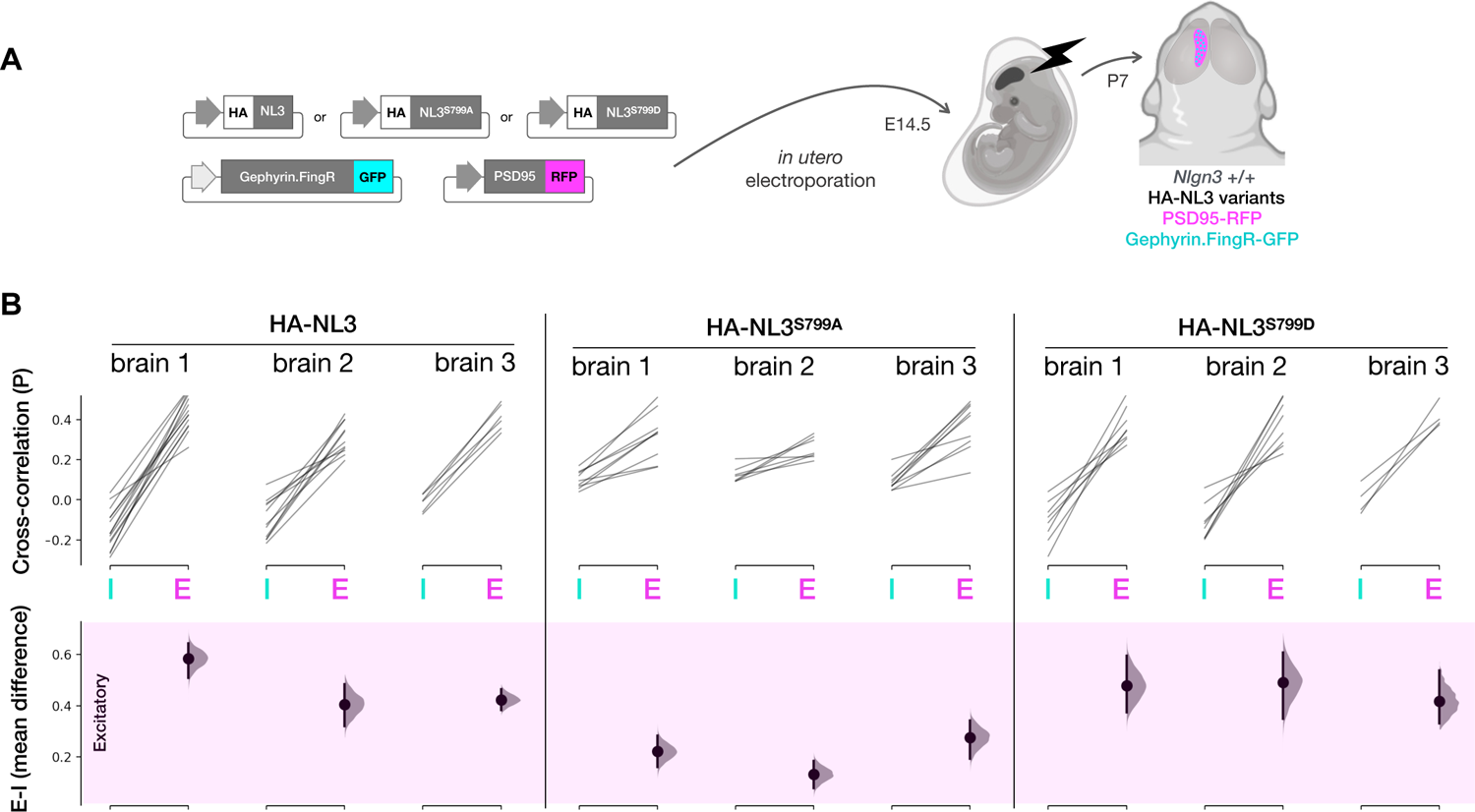
Phosphomimetic variants of HA-NL3 coexpressed with excitatory and inhibitory postsynapse markers in WT mouse cortex. (A) Schematic of *in utero* electroporation approach to express HA-tagged NL3 variants and postsynaptic markers in mouse cortex. Plasmid cocktails were used to test each HA-tagged NL3 variant (WT, S799A phospho-null, and S799D phosphomimetic) consisting of Gephyrin.FingR-GFP to label inhibitory postsynapses, and PSD95-RFP to label excitatory postsynapses. Plasmid cocktails were electroporated into E14.5 mouse embryos and brains were imaged at P7. (B) Specificity of excitatory versus Inhibitory postsynapse localization was quantified in three brains for each HA-NL3 variant on a WT background. Localization was quantified as in Fig. 5, based on cross-correlation analysis of the HA signal to GFP inhibitory (I) and RFP excitatory (E) synapse marker signals. Top plot shows average cross-correlation values (P) in paired E-I analyses from each brain section in each of three brains per variant. Bottom plot shows paired mean differences of excitatory versus inhibitory correlations per brain in Cumming estimation plots. Effect size is shown with sampling distributions and vertical bars indicating 95% confidence interval. Positive values indicate specificity of HA-NL3 variants for excitatory postsynapses, while negative values would indicate specificity of HA for inhibitory postsynapses. All HA-NL3 variants preferentially localize to excitatory synapses in cortex, possibly reflecting intrinsic dimerization of exogenous HA-NL3 phosphomimetic variants with endogenous WT NL3 as previously reported (Poulopoulos 2013). The degree of specificity for excitatory postsynapse localization is highest in WT (0.47 ± 0.057) and phosphomimetic S799D mutant HA-NL3 (0.46 ± 0.023), while decreased specificity for excitatory versus inhibitory postsynapses is observed with phospho-null S799A mutant HA-NL3 (0.2 ± 0.039). Values are E-I mean difference from n=4 to 14 sections (depending on electroporation field size) from each brain, with deviations in SEM for N=3 brains per group.

## METHODS

### Plasmids

NL3 construct variants were produced from an HA-tagged expression construct of NL3 previously cloned from rat cDNA into pCMV^19^. Site-directed mutagenesis using the Quikchange method (Stratagene) was used to produce S799A and S799D mutants. Full length HA-NL3 and respective mutants were subcloned into vector pRaichu for expression in cell lines or neurons. Cytosolic domains of NL3 and corresponding mutants were subcloned into vector pLexN for yeast-two-hybrid assays. pRaichu (generously provided by Jun-ichi Miyazaki, Osaka) is based on the pCAGGS vector backbone^65^. pLexN is prey construct modification of pBTM116^66^. pEGFPN1, pRaichu-NL3 and corresponding S799A and S799D variants, as well as pRaichu-NL3-ΔICD (NL3 with stop codon replacing position K735) were used for transfections. Previously published myc-CB2^SH3-67^ and GFP-Gephyrin^68^ were used for COS7 cell transfections. For *in utero* electroporation, NL3 and corresponding mutants were subcloned into pJac, made from pCX with the promoter replaced by human Synapsin (hSyn) using golden gate assembly (NEB). For synaptic labelling pCAG-GPHN-FingR-EGFP-CCR5TC^40^ Addgene # 46296) and pLL(Sy)-PSD95-TagRFP^39^ were used. For CRISPR KO, plasmid pJ2_SpCas9 was constructed using golden gate assembly to express spCas9 driven by a CAG promoter.

In order to create gRNA-resistant NL3 and corresponding mutants, overlap extension PCR (KAPA HiFi 2X RM) was carried out using primers:

Forward 1 ggtctcaaggtgccaccATGTGGCTGCAGCTCG

Reverse 1 GAGCACGGAGCCGTCGATCATATTTC

Forward 2 GAAATATGATCGACGGCTCCGTGCTC

Reverse 2 ggtctcaaagcCTATACACGGGTAGTGGAGTG.

For guide RNA targeting NL3, primers 5’ caccGGAACAGGCAACATGATTGA 3’ and 5’ aaacTCAATCATGTTGCCTGTTCC 3’ were annealed and golden gate assembled using BpiI (Thermo Fischer) and T4 Ligase (NEB) into s23_U6_scaffoldv2 backbone having BpiI cut sites downstream of U6 promoter.

For yeast-two-hybrid assays, the intracellular domains of rat NL1-NL3 and mouse NL4 were subcloned into pLexN to serve as bait constructs, as previously described (Poulopoulos 2009). Corresponding mutant bait constructs of NL3 S799A and S799D as well as the homologous mutant variants of NL1, NL2, and NL4 were produced with site-directed mutagenesis. Plasmid vector pVP16-3^66^ was used for prey constructs of full-length Gephyrin^19^ and the sequence comprising the three tandem PDZ domains of S-SCAM (PDZ1-3, residues 422–497 of P_446073)^51^ Coding regions of all plasmids were verified by Sanger sequencing. Plasmid sources and sequences for custom-made plasmids are provided in the Reagent and Resource Table below.

### Animals

All animal experiments were designed and carried out in compliance with animal welfare guidelines of the European Community Council Directive 86/609/EEC and approved by the Bioethical Committee of Turin University, the Institutional Animal Care and Use Committee of University of Maryland School of Medicine, and by the state of Lower Saxony, Germany according to the corresponding permits 33.9-42502-04-13/1359, and 33.19-42502-04-13/1052. For information on mouse and rat experimental strains see Reagent and Resource Table below.

### Cell culture and transfection

COS7 cells were plated directly onto glass coverslips, cultured in DMEM supplemented with 10% fetal bovine serum, and transfected with FuGENE6 (Roche) according to standard protocols.

Primary neuron cultures were prepared from cortex of wild-type (WT) C57BJ embryos at embryonic day (E) 16. Cultures were prepared as described previously^69^, with papain substituted for trypsin to obtain better cell recovery following tissue digestion and trituration. Cells were plated on poly-L-lysine-coated glass coverslips at a density of 25,000-75,000 cells/cm^2^. Neurons were transfected at day in vitro (DIV) 3 with Lipofectamine 2000 using standard protocols. Analyses were carried out at DIV 7-14.

For CRISPR reagent validation, Neuro-2A cells were transfected with DNA plasmids expressing Cas9, GFP and NL3 guide RNA or scrambled guide RNA using transient transfection in 24 well plates. Briefly, cells were transfected at 70-80% confluence using Polyethylenimine, Linear, MW 25000 (‘PEI’, Polysciences) resuspended to 1mg/mL in H2O at a 3:1 (v/w) PEI:DNA ratio with 250 ng DNA per plasmid (750 ng total DNA) diluted in Opti-MEM (ThermoFisher Scientific) and added dropwise to cells. After 48 hours of transfection, cells were split into 6-well plates and allowed to grow for 5 more days.

### Cleavage assessment and sequencing of CRISPR editing outcomes

Mouse Neuro-2A cells were transfected with Cas9 and gRNA constructs targeting candidate targets on *Nlgn3*. Empty gRNA scaffold was used as control. At 7 days post transfection, genomic DNA was isolated using the Genomic DNA Clean & Concentrator kit (Zymo Research) and used as template for PCR using KAPA HiFi HotStart DNA Polymerase with 2X Master Mix (Roche) to amplify the target sequence in exon 4 of *Nlgn3* using forward 5’ CCCCAGAAGCTAGCCATGGTCAC 3’ and reverse 5’ GACAGGACCTATAAAACTCAGCCAAG 3’ primers. Batch amplicons were gel extracted with Zymoclean Gel DNA Recovery kit (Zymo Research), and Sanger sequenced (Genewiz) using primer 5’ GGTGGTGTCCTGGCACCTG 3’ to yield mixed sequence chromatographs representing all editing outcomes. Sequences were aligned using Benchling (https://benchling.com) and unmixed using Tracking of Indels by DEcomposition^70^ web tool (http://shinyapps.datacurators.nl/tide/) to quantify indels.

Split GFP recombination assays^46^ were used to assess cleavage efficiency of *Nlgn3* gRNAs. gRNA target region was cloned into pCAG-mTagBFP-2A-EGXXFP vector using Esp3I (Thermo Fischer) and T4 Ligase (NEB) using oligo annealing of 5’ctagTTCCTCCTATGTAGAGAGCCGTCACGAAT3’ and 5’aattATTCGTGACGGCTCTCTACATAGGAGGAA3’. HEK 293T cells were transfected with DNA plasmid of split GFP reporter and plasmids expressing RFP_2A_Cas9 and gRNA::*Nlgn3* or control gRNA. Percentage of GFP positive cells over BFP (split GFP plasmid reporter) and RFP (Cas9 reporter) expressing cells were analyzed using General Analysis tool (NIS Elements) in control and gRNA:::*Nlgn3* groups.

### Antibodies

The following antibodies were used for immunocytochemistry on cell lines and cultured neurons (ICC), immunohistochemistry on brain sections (IHC), immunohistochemistry on brain sections after *in utero* electroporation (IUE), Western blotting (WB), and immunoprecipitation (IP): Mouse-anti Actin (AC40, Sigma, 1:5000 for WB), Mouse-anti-myc (9E10, 1:1,000, Sigma-Aldrich for ICC), Mouse anti-gephyrin (3B11; Synaptic Systems; 1:500 dilution for ICC, IUE); mouse anti-gephyrin (mAb7a; Synaptic Systems; 1:1000 dilution for IHC); mouse anti-PSD95 (6G6-1C9; Abcam; 1:500 dilution for ICC); mouse anti-PSD95 (clone K28/43; NeuroMab; 1:1000 dilution for IHC); rabbit anti-GFP (598 MBL; 1:1000 dilution IUE); rabbit anti-GFP (132003; Synaptic Systems; 1:1000 dilution for IC); guinea pig anti-GABA_A_Rγ2 (1:2000; provided by Dr. Jean-Marc Fritschy, Zurich); mouse anti-GlyRα1 (Mab4a; 1:100; provided by Dr. Heinrich Betz, Frankfurt); guinea pig anti-VIAAT (131004; Synaptic Systems; 1:1000 dilution for IC, IUE); guinea pig anti-VGluT1 (135304; Synaptic Systems; 1:1000 dilution for IC, IUE); rabbit anti-HA (715500; Invitrogen; 1:1000 dilution for IC, 1:4000 dilution for WB).

Custom-made NL3 antibody 804 (1:1000 dilution for WB, 5-10 μl antiserum for IP) was previously described and validated in NL3 KO samples^34^, and by mass spectrometric identification of NL3 in the material immunopurified using 804 antibody from rat brain lysate (Fig. 2). Custom-made phospho-specific antibody 6808 recognizing the serine-phosphorylated NL3 epitope RRpSPDDIP, was produced by PhosphoSolutions (Aurora, CO, USA). Briefly, antiserum was raised in rabbits immunized with RRpSPDDIP. Antibody was affinity-purified against the immunizing phosphopeptide. This produced antibody RαNL3-pS 6808, which recognizes NL3 phosphorylated at S799, and is likely to recognize any putative phosphorylation at the homologous Gephyrin-binding site serine of NL1 or NL4. 6808 was used for WB at a dilution of 1:3000 in the presence of 2 μg/ml of competing unphosphorylated peptide RRSPDDIP. Peptides were synthesized in-house (Neuroproteomics Group, MPI-NAT) by standard solid phase peptide synthesis using Fmoc chemistry.

The following secondary antibody conjugates where used for ICC, ICH, and IUE (listed by host and dye): goat Alexa 488, donkey Alexa 488, goat Alexa 568, donkey Alexa 568 (Molecular Probes, Eugene, OR); goat Cy3, donkey Cy3, goat Cy5, donkey Cy5 (Jackson Immunoresearch, West Grove, PA). Secondary antibodies conjugated with HRP (Jackson ImmunoResearch) or IRDye800 (Rockland) were used for WB. For details and antibody sources see Reagent and Resource Table below.

### Immunopurification and Mass Spectrometry (MS)

One adult rat brain (2 g) was homogenized in nine volumes of 320 mM sucrose solution supplemented with cOmplete Protease Inhibitor Cocktail (Roche) and Phosphatase Inhibitor Cocktail 1 and 2 (Sigma) at manufacturer recommended concentrations. Homogenates were produced at 4°C with 13 strokes in a Potter-Elvehjem homogenizer rotating at 900 rpm. Homogenates were centrifuged at 1200 x g for 15 minutes to obtain the supernatant postnuclear homogenate. Membranes were pelleted from the postnuclear homogenate with 100,000 x g for 1h. Membrane proteins were extracted from pellets with 1% SDS in TNE buffer (50 mM Tris, 150 mM NaCl, 5 mM EDTA). SDS lysate was centrifuged at 20,000 x g for 30 minutes and supernatants were subsequently diluted with 7 volumes of 1.15% Triton-X 100 in TNE with Phosphatase Inhibitor Cocktail 1 and 2 (Sigma) to produce a mixed-micelle lysate with final detergent concentrations of 0.125% SDS and 1% Triton-X 100, which allow immunopurification procedures (Poulopoulos 2009). Endogenous NL3 was immunopurified from this lysate by using the anti-NL3 antibody 804, preloaded onto Protein G Sepharose beads (Amersham). After overnight incubation at 4°C with the lysate, beads were washed three times with TNE buffer containing 1% Triton-X 100 and once with TNE buffer alone. Proteins bound to beads were eluted in SDS sample buffer, separated by SDS-PAGE on precast 10% Bis-Tris gels (NuPAGE, Thermo Fisher Scientific) with MOPS running buffer, and visualized by colloidal Coomassie staining.

Gel bands of interest were excised and subjected to in-gel digestion with endoproteinase Asp-N to generate a proteolytic peptide that covers the potential phosphorylation sites within the Gephyrin-binding site of NL3 and is of a size readily accessible for mass spectrometric sequencing (which is not possible with trypsin because of the lack of cleavage sites within a ∼40 amino acid sequence stretch C-terminally of the Gephyrin-binding site, Fig. 2A). Phosphopeptides were enriched by TiO_2_ chromatography as described (Oellerich 2009) and the presence of a monophosphorylated species of the peptide NL3 (residues 791-800, DYTLTLRRSP, M_calc_ = 1300.62) in the enriched fraction was confirmed by MS (data not shown). This target phosphopeptide was sequenced by NanoLC-ESI MS analysis on an Ultima API-Q-TOF mass spectrometer (Waters Cooperation) as described (Oellerich 2009).

### Yeast-two-hybrid

Yeast-two-hybrid (YTH) assays were performed in the *Saccharomyces cerevisiae* L40 reporter strain using small-scale Li-acetate cotransformations with pLexN bait constructs encoding the intracellular domains of NL1-NL4 and either empty pVP16-3 or pVP16-3 vectors encoding full-length Gephyrin or S-SCAM fragment PDZ 1-3 (residues 422–497). Transformed yeast were incubated at 30°C for three days to allow prototropic colonies to emerge as described previously^71^. Clones were tested in duplicate for activation of the β-galactosidase reporter gene by filter assays^66^. The readout corresponds to LacZ activity using standard X-gal chromogenic assay.

### In vitro kinase assays

For Kinase Hot-Spot assays, a 21 amino acid peptide spanning residues 785-805 of rat NL3 was used as substrate for *in vitro* kinase assays. Streamlined filtration binding assay was performed on a commercial basis by the Reaction Biology Corporation (PA, USA) in buffer containing 20 mM HEPES (pH 7.5), 10 mM MgCl_2_, 1 mM EGTA, 0.02% Brij35, 0.02 mg/ml BSA, 0.1 mM Na_3_VO_4_, 2 mM DTT, 1% DMSO in the presence of ATP-γ-^33^P and substrate peptide, as previously^72^. Experiments were carried out in single dose duplicates with results expressed as percent incorporation relative to internal control, or as nM phosphate transferred to 10 μM peptide (see Extended Data Fig. 3). ADP Glo assays (Promega) were performed according to manufacturer’s recommendations. WT rat NL3 peptide representing residues 785-805, and the three mutant variants Y793F, T795A, and S799A were examined. All peptides were synthesized in-house (Proteomics Group, MPI-EM) by standard solid phase peptide synthesis using Fmoc chemistry.

### Western blot

Adult mouse brains from WT and KO littermates were dissected into regions of interest and manually homogenized in Lysis Buffer (0.32 M Sucrose, 5 mM MgCl_2_ supplemented with protease inhibitor cocktail, phosphatase inhibitor cocktails, 200 U/mL benzonase) using a Teflon pestle in microcentrifuge tubes. Total protein concentrations were determined using Bio-Rad protein assay according to the manufacturer’s instructions. Homogenate samples were prepared in Laemmli Buffer (10% Glycerol, 50 mM Tris-HCl pH 6.8, 2 mM EDTA, 2% SDS, 100 mM DTT, 0.05% Bromophenol blue) at 65 °C for 20 min. Samples corresponding to 40 μg total protein were loaded for SDS-PAGE. Proteins separated on gels were electroblotted onto nitrocellulose (Protran 0.2 μm, GE Healthcare) with a constant current of 100 mA for 10 h using wet transfer. For Immunoblot, membranes were incubated in Blocking Buffer (5% BSA in TBS-T) for 1 h at RT. Standard western blot procedures were followed with overnight incubations of primary antibody solutions in Blocking Buffer at 4 °C. For NL3-pS WB, primary antibody 6808 solution was pre-incubated with 2 mg/mL unphosphorylated antigen peptide (RRSPDDIP) for 1h at RT prior to primary antibody incubation. HRP or IRDye800 conjugated secondary antibodies were incubated with membranes for 1h at RT in Blocking Buffer. Immunoblot signal was detected using enhanced chemiluminescence (ECL, GE Healthcare) or with the Odyssey Infrared Imaging System (LI-COR Biosciences), respectively. Experiments were repeated and confirmed using WT rat brain as well.

### Spot blot

Peptide stock solutions containing 2 mg/mL of NL3 antigenic peptides (unphosphorylated RRSPDDIP or phosphorylated RRpSPDDIP) were prepared in TBS (20 mM Tris-HCl pH 7.5, 137 mM NaCl). Peptide stock solutions were serially diluted into 1000 ng/µL, 250 ng/µL, 100 ng/µL and 50 ng/µL. 1 µL of each serial dilution of phosphorylated and non-phosphorylated peptides was spotted onto nitrocellulose membranes. Immunoblot against NL3-pS was performed using the protocol described above for western blots.

### Membrane co-clustering assay

Membrane co-clustering assays were performed in COS7 cells transfected with GFP Gephyrin, myc-CB2^SH3-^ and HA-NL3 variants as described previously^19^. Briefly, cells were fixed 10 h after transfection in 4% PFA and blocked with 5% normal goat serum and 0.1% gelatin in 0.1 M phosphate buffer (PB). Prior to permeabilization, cells were stained with polyclonal rabbit anti-HA antibody in blocking solution for 2 h at RT to detect the surface pool of HA-tagged NL3. After three washes with PB, cells were permeabilized with 0.1% Triton X-100, 5% normal goat serum, and 0.1% gelatin in PB, and stained with mouse-anti-myc antibody (clone 9E10, 1:1,000, Sigma-Aldrich) in the same buffer for 2 h at RT, followed by the appropriate Alexa-conjugated secondary antibodies.

Samples were imaged using a Leica DMIRE2 microscope equipped with a 63× oil-immersion objective connected to a Leica TCS SP2 AOBS confocal laser scanning setup. Intensity correlation analysis was performed on multichannel images using ImageJ. Briefly, a Gaussian blur was applied, and GFP and HA channels were thresholded. A standard Pearson’s correlation coefficient was evaluated between the HA and GFP channels in the thresholded fields using the Intensity Correlation Analysis plugin for ImageJ (http://rsb.info.nih.gov/ij/) from T. Collins and E. Stanley (Toronto, ON, Canada).

### Synaptic and morphological analyses in cultured neurons

Quantification of synaptic puncta and neuron morphology were carried out in cultured mouse cortical neurons at DIV7-14 transfected with HA-NL3 variants and stained for HA and endogenous synaptic markers. Images for pre- or postsynaptic quantifications were captured and analyzed in a double-blind manner wherever possible. For quantification of presynaptic VGluT1, VIAAT, and postsynaptic Gephyrin or PSD95, each image was manually thresholded in ImageJ to generate a binary image in which individual clusters of synaptic molecule markers could be observed. In order to isolate puncta of transfected neurons only, a mask was created from a co-stained image captured for HA. This mask allowed quantification of synaptic puncta selectively in transfected neurons. The number of clusters per neuron was counted using the ‘Analyze Particles’ function with Watershed Segmentation. The total mask area was measured with ‘measure’ function. Puncta were expressed as number of puncta per given area. For DIV7 cultures where synaptic puncta have not fully matured, we assessed total intensity of Gephyrin and PSD95 within the HA mask of transfected neurons. The intensity was normalized to total image intensity. Results were expressed as means of four independent experiments.

Neuronal arborization was assessed by thresholding the HA-stained image, creating a binary image in which all neuronal branches are visible and distinguishable. Axons were manually removed using selection tools, as were any other dendrites or axons not originating from the selected neuron. Sholl analysis (Sholl, 1953) was carried out with a radius of 10 μm to 100 μm from the soma. The number of intersections was plotted against distance from the soma. Analysis of filopodia was performed by manual counting of filopodial protrusions along 10 μm segments of 2^nd^ and 3^rd^ order dendrites. Three dendrites were assessed per image with at least 15 images per condition.

### Immunohistochemistry on brain sections

Adult mice were anesthetized with intraperitoneal ketamine-xylazine 1:1 (0.1 ml/kg) and decapitated. Brains were dissected out and cut into sagittal or coronal slabs of ≈1 mm and fixed by immersion in ice-cold formaldehyde (4% in 0.1 M phosphate buffer) for 20–30 min as described previously^73, 74^. The slabs were cryoprotected in ascending sucrose solutions (10%, 20%, and 30%) and sectioned with a cryostat. Following a blocking step in normal goat (or donkey) serum (3% in PBS with 0.5% Triton X-100), sections were incubated overnight with combinations of two or three primary antibodies raised in different species. The sections were then rinsed in PBS, incubated with appropriate secondary antibodies, rinsed again and coverslipped with Dako fluorescence mounting medium (Dako Italia, Italy).

Immunolabeled sections were imaged using a laser scanning confocal microscope (Zeiss LSM5 Pascal) using the multichannel acquisition mode to avoid fluorescence crosstalk. Images (512 × 512 pixels) were acquired with a 100× oil-immersion objective (1.4 numerical aperture) at a magnification of 8.1 x 10^-3^ or 2 x 10^-3^ μm^2^/pixel (zoom 2 and zoom 4, respectively), and the pinhole set at 1 Airy unit.

Acquired images were processed with the image-analysis program Imaris (release 4.2; Bitplane, Zurich, Switzerland). To analyze colocalization between NL3 and gephyrin or PSD95, images were first segmented using a threshold that maximized the selection of immunofluorescent puncta over background labeling, and then processed with the “colocalization” module, in which a mask is generated from the comparison of two different confocal channels. The number of puncta was then calculated with NIH ImageJ software (http://rsb.info.nih.gov/nih-image) as previously described^74^.

For analysis of in utero electroporated brains (Fig. 5 and Extended Data Fig. 6), anesthetized electroporated animals were perfused first with ice-cold PBS and then with ice-cold formaldehyde (4% in 0.1 M phosphate buffer). Brains were dissected out and coronal slices (50-80 um) were obtained using a Leica vibratome VT1000S. Staining were performed using a blocking solution 5 % BSA, 0.3% Triton X-100 in PBS. After blocking step, primary antibodies were incubated overnight at room temperature. The sections later were rinsed with PBS and incubated with appropriate secondary antibodies for 5h at room temperature. Lastly, slices were rinsed with PBS and coverslipped using Fluoromount-G with DAPI (Invitrogen #00-4959-52).

Immunolabelled sections were imaged using a laser scanning confocal microscope (Nikon A1) with 60X oil immersion objective with 1.55 digital zoom using the multichannel acquisition mode to avoid fluorescent crosstalk. Acquired images were processed with NIS Elements analysis software (refer to Quantification and Statistical Analysis).

### In Utero Electroporation

In utero electroporation of plasmid DNA were performed on embryonic day 14.5 (E14.5) to target cortical layer II/III, as previously described^75, 76^. Briefly, DNA solutions were prepared to 4 μg/μL total DNA, with equal amount of each plasmid (HA-NL3; PSD95-tagRFP; Gephrin.FingR-GFP for WT background experiment or HA-NL3; PSD95-tagRFP; Gephrin.FingR-GFP; Cas9; gRNA for CRISPR KO background experiment). Dames were deeply anesthetized with isoflurane under a vaporizer with thermal support (Patterson Scientific Link7 & Heat Therapy Pump HTP-1500). After abdominal area preparation by hair removal and surgery scrub (70% ethanol and 10% Betadine solution), laparotomy is carried out to expose the uterine horns and embryos. Using a glass micropipette pulled (Narishige PC-100) and beveled (Narishige EG-45), DNA solution was injected into lateral brain ventricles, followed by application of 4 x 50 ms square pulses of 35 V (NEPA21 electro-kinetic platinum tweezertrodes connected to a BTX ECM-830 electroporator) to target cingulate cortex. Typically, 4-6 pups were electroporated per dame. Later, uterine horns and embryos were placed back inside the abdominal cavity and muscle and skin incisions were closed by monofilament nylon sutures (AngioTech). After term birth, electroporated mouse pups were non-invasively screened at the age of P0 for cortical fluorescence using a fluorescence stereoscope (Leica MZ10f with X-Cite FIRE LED light source) and returned to their dame.

### Quantification and Statistical Analysis

Puncta were defined by intensity thresholding of the immunoreactive signals, and the proportion of overlapping puncta was determined as previously described^74^. Due to the different densities with which PSD95 and Gephyrin puncta present in the brain, we followed distinct approaches to quantify the degree and specificity of overlap in each case. PSD95 displays high punctum densities covering much of grey matter neuropil. As such, colocalization alone is a poor readout of specific association due to the high incidence of random overlap. Thus, random overlap with PSD95 immunoreactivity was assessed and compared to specific overlap.

For analysis of in utero electroporated brains (Fig. 5 and Extended Data Fig. 6), puncta were defined by intensity, size, and circularity of the signal. In order to analyze colocalization of NL3 and Gephyrin or PSD95, GA3 Analysis (NIS Elements) workflow has been created. Briefly, Gephyrin and PSD95 puncta were segmented using a threshold function (using intensity, size, and circularity parameters) to obtain very precise selection of puncta for each channel. Later, binary calculation was performed to combine PSD95 and Gephyrin puncta as double binary. Pearson correlation coefficiency was used to analyze colocalization between Gephrin-NL3 and PSD95-NL3 on double binaries.

Statistical comparisons in this study were made using the unpaired, two-tailed Student’s *t*-test when comparing two variable means. In the case of three variables, we used the one-way ANOVA with Bonferroni posthoc test. *p* values < 0.05 were considered significant. p value outcomes were indicated by asterisks in the figures as follows: *\*p*<0.05, *\*\*p*<0.01, and *\*\*\*p*<0.001.

### Reagent and Resource Table

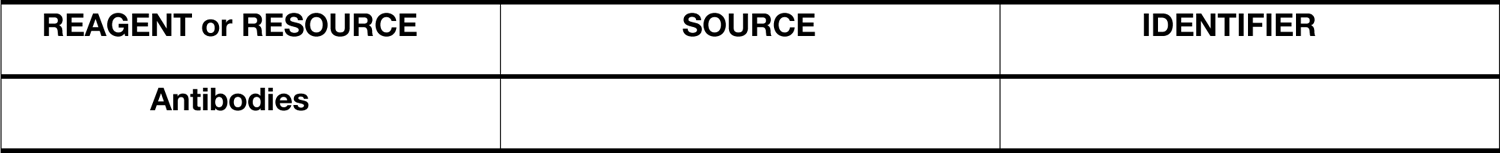
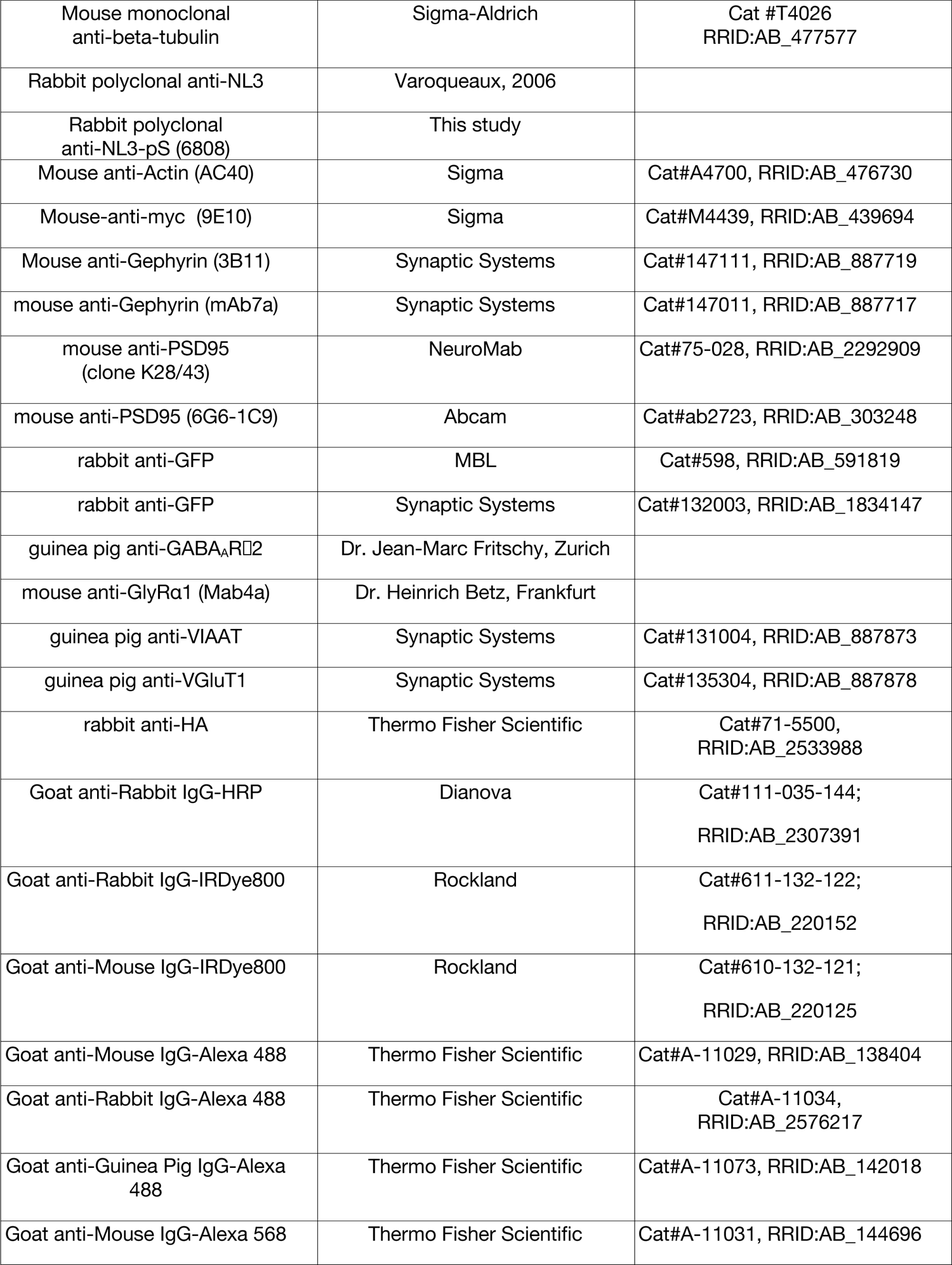
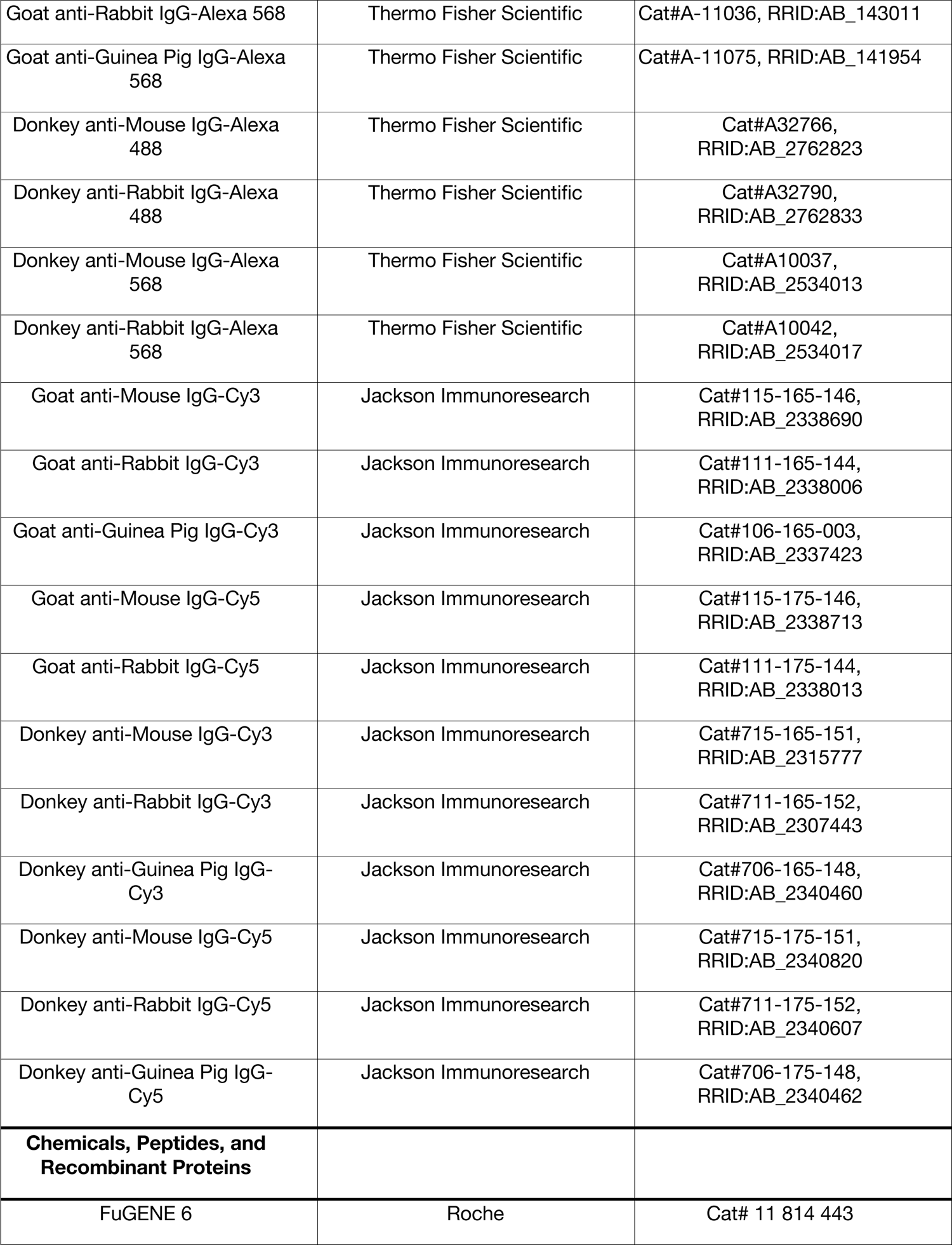
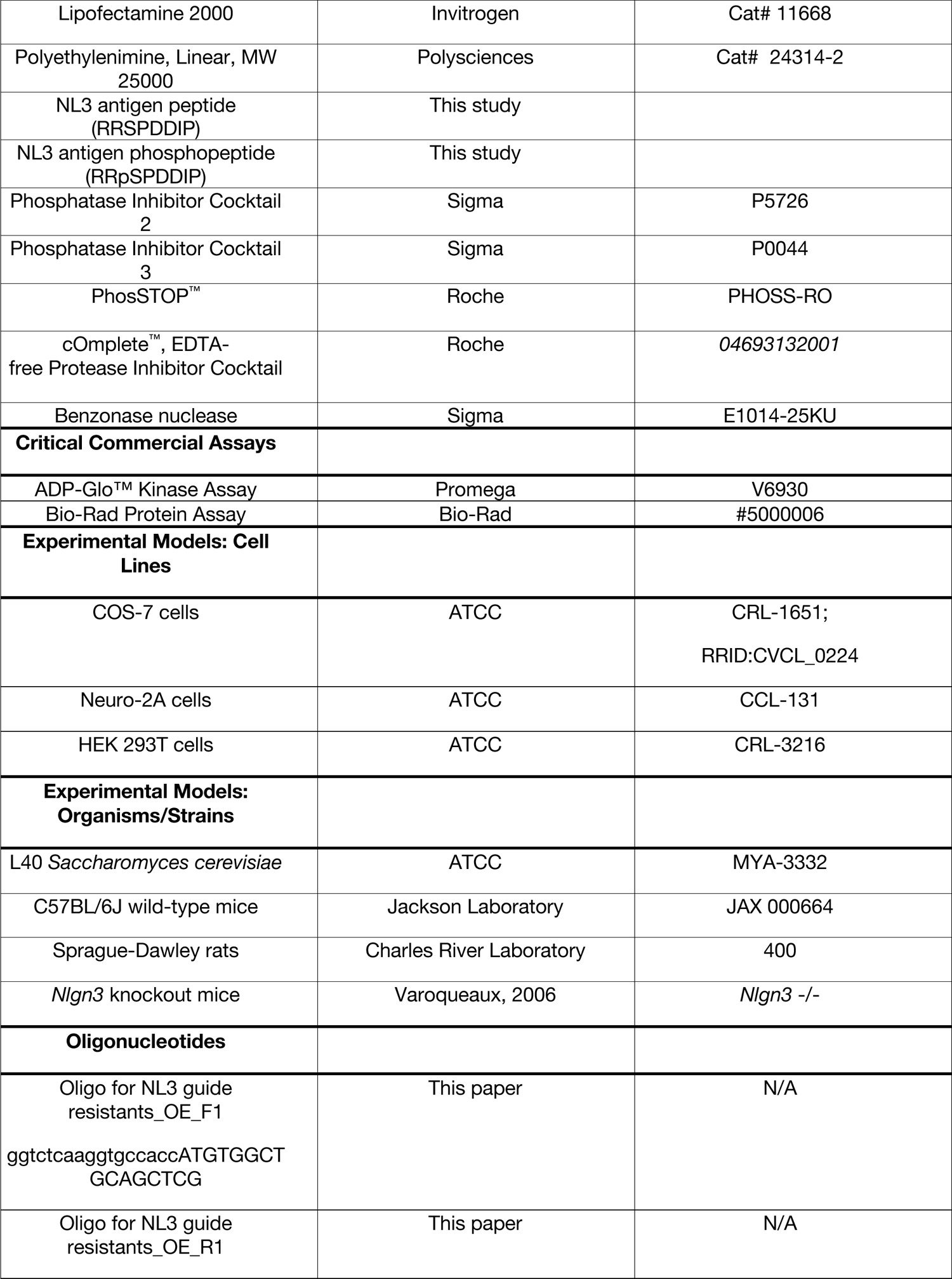
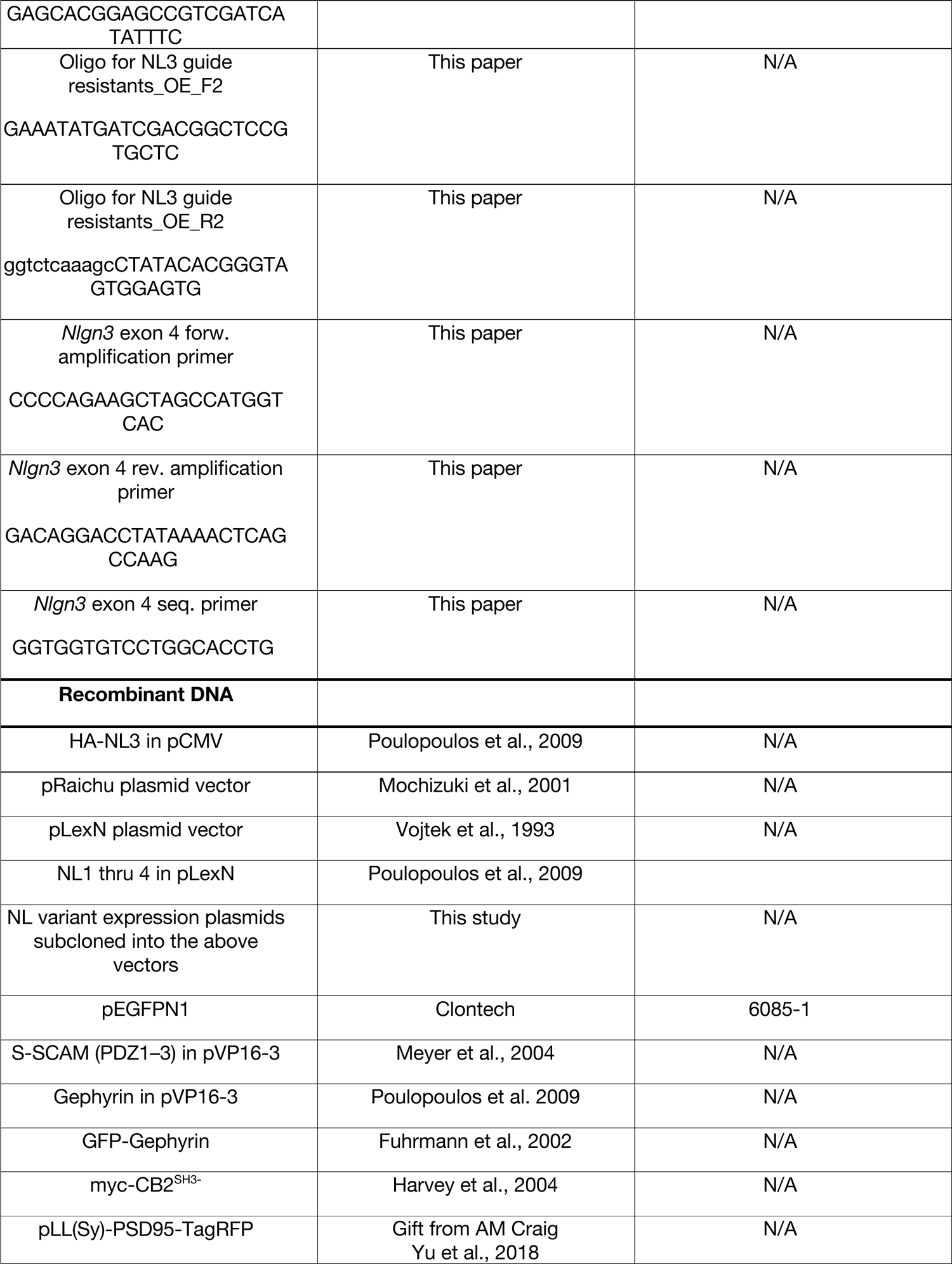
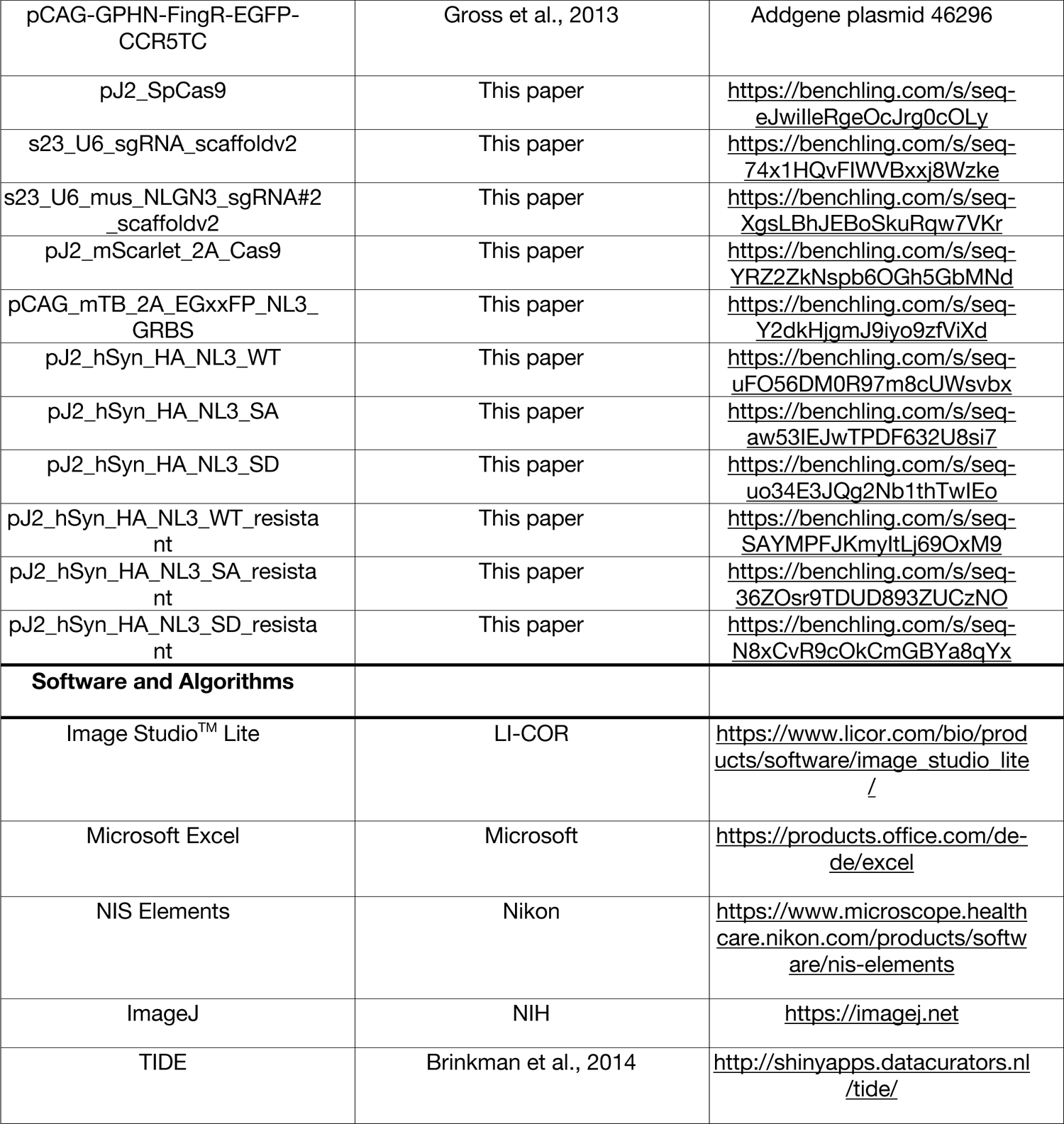

